# Structural and mechanistic insights into the DNA glycosylase AAG-mediated base excision in nucleosome

**DOI:** 10.1101/2022.06.16.496441

**Authors:** Lvqin Zheng, Bin Tsai, Ning Gao

## Abstract

DNA glycosylase engaging with damaged base marks the initiation of base excision repair. Nucleosome-based packaging of eukaryotic genome obstructs DNA accessibility, and how DNA glycosylases locate the substrate site on nucleosomes is currently unclear. Here, we report cryo-electron microscopy structures of nucleosomes bearing a deoxyinosine (DI) in various geometric positions and structures of them in complex with DNA glycosylase AAG. The apo nucleosome structures show that the presence of a deoxyinosine alone perturbs nucleosomal DNA globally, leading to a general weakening of the interface between DNA and the histone core and a greater flexibility to the exit/entry of the nucleosomal DNA. AAG makes use of this nucleosomal plasticity and imposes further local deformation of the DNA through the formation of the stable enzyme-substrate complex. Mechanistically, local distortion augment, translation/rotational register shift and partial opening of the nucleosome are employed by AAG to cope with substrate sites in fully exposed, occluded and complete buried positions, respectively. Our findings reveal the molecular basis for the DI-induced modification on the structural dynamics of the nucleosome and elucidate how DNA glycosylase AAG accesses damaged sites on the nucleosome with different solution accessibility.

## Introduction

The eukaryotic base excision repair (BER) machinery locates and repairs DNA base damage in chromatin(*1, 2*). Genomic DNA contains substantial amounts of DNA base damages due to various exogenous damaging agents and spontaneous decomposition reactions—such as deamination of deoxycytidine (DC, cytosine) to deoxyuridine (DU, uracil) and deamination of deoxyadenosine (DA, adenine) to deoxyinosine (DI, hypoxanthine)(*3*). In human cells, DNA base damage is primarily detected by damage-specific DNA glycosylases(*4, 5*). DNA glycosylase recognizes damaged base and catalyzes its excision, leaving an apurinic/apyrimidinic site (AP site) which will be cleaved by AP site endonuclease APE1 to generate a nick on DNA (*6*). Subsequent repair reactions involve AP-lyase activity, DNA polymerase activity and DNA ligase activity, these reactions are executed by XRCC1, DNA polymerase β and DNA Ligase III in the short-patch BER sub-pathway, while the long-patch BER sub-pathway requires the replicative DNA polymerase δ/ε-PCNA-RFC machinery, flap endonuclease 1 and DNA ligase I(*7*).

DNA glycosylase AAG as one of the first responders of DNA base damage can recognize alkylpurines like 3-methyladenine and 7-methylguanine(*8, 9*), oxidized adenine 1,N^6^-ethenoadenine(*10*) and deaminated adenine hypoxanthine (*6, 11–13*). In humans, altered expression and single nucleotide polymorphisms of AAG are associated with microsatellite instability(*14*), spontaneous frameshift mutagenesis(*15*), and a variety of cancers including osteosarcoma(*16*), breast cancer(*17, 18*) and astrocytic tumors(*19, 20*). In mouse models, Aag-knockout mice is prone to liver cancer, while excessive AAG activity in Aag-overexpressing mice causes hepatotoxicity and lethality(*21*).

Frequently occurring DNA base damages could be present in all regions of chromatinized eukaryotic genome, including nucleosomal DNA sites where nucleosomes would obstruct base excision repair machinery’s accessibility to the damage sites(*22*). In a nucleosome, the 147-bp DNA wraps around the octameric core in 1.65 superhelical turns, leaving only a portion of solvent-facing DNA freely accessible(*23, 24*). Extensive *in vitro* studies showed that AAG (*25, 26*) along with short-patch BER factors(*27–29*) could selectively locate and bind to the damage sites in nucleosomes. In general, AAG activity on nucleosome is strongly correlated to solution accessibility of damaged base, and *in vivo* studies also confirmed this general correlation (*30*), and found that bases at the DNA minor grooves, which interact with the octameric histone core, display higher mutation rates due to low accessibility to repair factors(*22*). Structurally, the solution accessibility is predominantly determined by its translational position and rotational orientation on a nucleosome, and solvent-facing damaged bases are indeed excised more efficiently than occluded and embedded damaged bases with medium and low solution accessibility (*31*). Another contributing factor is the structural dynamics of nucleosome such as spontaneous opening of nucleosome (*32, 33*), which could impact on the accessibility of the damaged bases on certain positions. In addition, compared to the other regions of the nucleosomal DNA, the dyad axis is the least accessible to repair factors (*31, 34*). Overall, the differential AAG activity at various nucleosomal positions could not be solely explained by the solution accessibility predicted from a static nucleosome structure.

While the structural mechanism of base excision enzymes’ action on naked DNA duplex is well-understood (*4, 35–40*), how these proteins overcome nucleosome-imposed obstacle to locate and repair DNA base damage in nucleosome is not fully understood. In the present study, we employed cryo-EM to explain how the base excision repair factor AAG accesses a damaged base introduced to various positions on nucleosomes. Our results show that a single DI nucleotide could induce a global perturbation to the nucleosome structure, and AAG exploits altered nucleosome dynamics and adopts distinct mechanisms to access the substrate site dependent on the geometric positions of the damaged base on the nucleosome.

## Results

### Preparation and structural determination of the AAG-NCP complexes

To capture the state of DNA glycosylase AAG engaging with DNA base damage in nucleosome, we first assembled nucleosome core particles (NCP) from a 152-bp Widom 601 DNA and core histones H2A, H2B, H3 and H4 from *Xenopus laevis* (Figure S1A)(*41–43*). The bottom strand of Widom 601 DNA (designated as damaged strand) contains a DI to mimic a base damage resulting from deamination of DA (Figure 1A).

**Figure 1.**
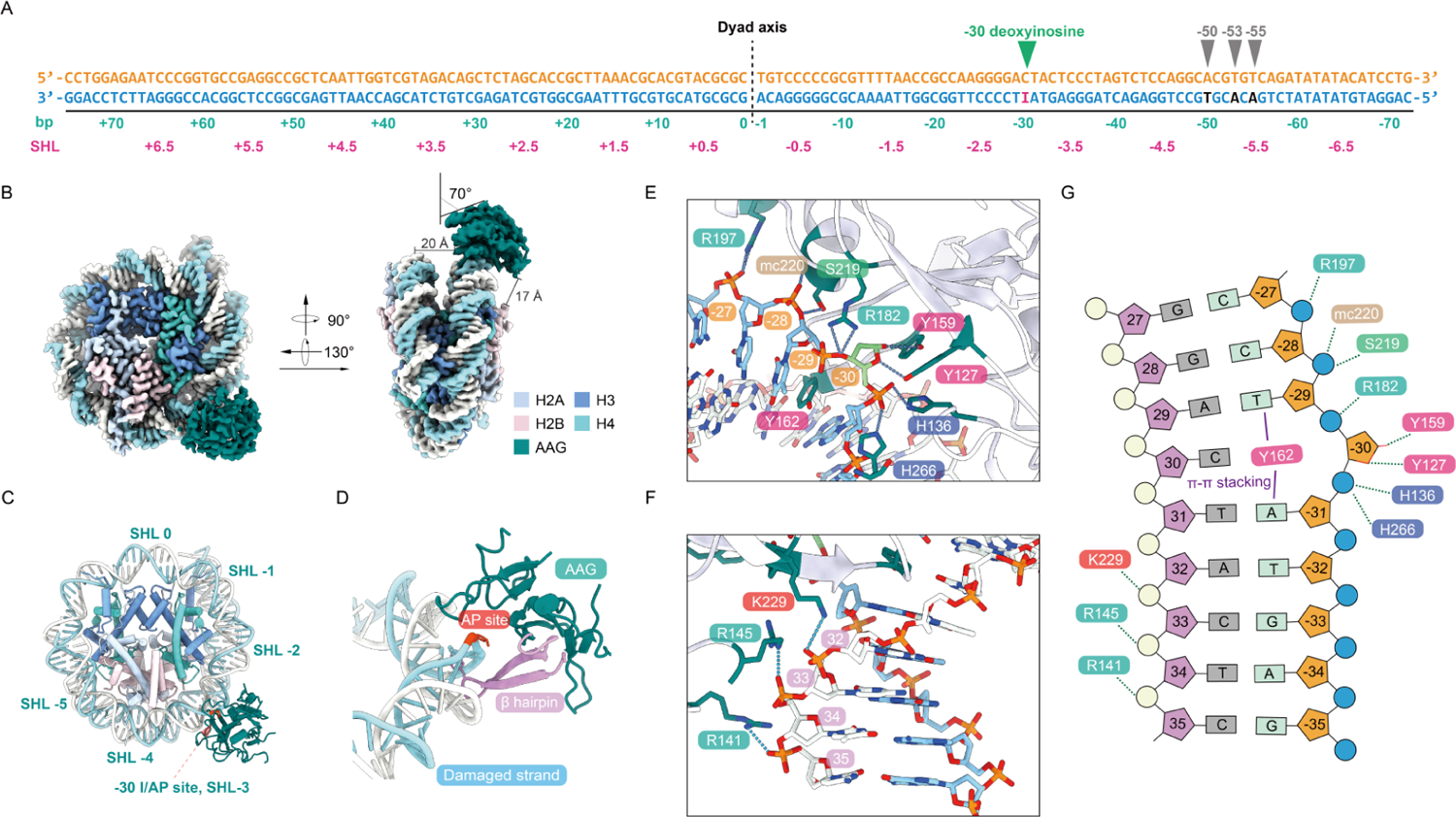
Cryo-EM structure of AAG-NCP^-30AP^ complex. **A**, Widom 601 sequence bearing deoxyinosine in various positions (−30, −50, −53 and − 55). **B**, Cryo-EM structure of DNA glycosylase AAG in complex with an NCP bearing deoxyinosine/AP site at −30 position, designated as AAG-NCP^-30AP^ complex. **C**, Atomic model of AAG-NCP^-30AP^ complex. **D**, A protruding β hairpin (β3-β4) of AAG is inserted into the minor groove of the nucleosomal DNA. **E**, Close-up view of the contacts between AAG and the deoxyinosine/AP site containing damaged strand of the nucleosomal DNA. **F**, Close-up view of the contacts between AAG and the undamaged strand of the nucleosomal DNA. **G**, Schematic representation of the atomic contacts in AAG-NCP^-30AP^ complex.

To investigate how AAG acts in positions with different solution accessibility and histone microenvironment, we sampled different superhelical locations (SHLs) and rotational orientations on nucleosome, a series of single DI-containing DNAs with a DI located in various geometric positions (−30, −50, −53 and −55) were prepared (Figure 1A). −30 at SHL-3 and −50 at SHL-5 both represent solvent-facing position with high solution accessibility, while −53 and −55 at SHL-5 represent occluded and imbedded position with medium and low solution accessibility, respectively(*31*). We assembled these NCPs and incubated them with purified AAG to form AAG-NCP complexes, which were further analyzed by cryo-EM (Figures S2-S5).

### Overall structure of the AAG-NCP^-30AP^ complex with a fully exposed damage site

The presence of DI in the −30 position of nucleosomal DNA (NCP^-30I^), which is solvent facing, triggers a strong engagement of AAG, as shown in the glycerol density gradient centrifugation (Figures S1B-S1C). Cryo-EM analysis of the AAG-NCP^-30I^ sample obtained high-resolution structures for the NCP in both AAG-bound and unbound states, at resolutions of 2.9 and 2.8 Å, respectively (Figures S2 and S6-S7). In the map of the free NCP (NCP^-30I^), the DI in the −30 position is well resolved and forms a non-canonical base pair with the DC on the top strand (Figure S6A), reflecting a state for the damaged nucleosome before AAG binding. In contrast, the map of the AAG-bound NCP was identified as a post-catalytic state (AAG-NCP^-30AP^). The ribose ring of the −30 position is in a flipped-out orientation, and there is no density for hypoxanthine base at −30, indicating that this is a post-catalytic AP site (Figure S6B). In addition, superimposition of a DI onto the catalytic cave comprising residues Y159, Y127, L180, R182, E125 and C167 would result in multiple steric clashes (Figure S6C).

This high-resolution structure of the AAG-NCP^-30AP^ complex provides an accurate model explaining the molecular details of the interactions between the damaged NCP and AAG. In general, AAG attaches to the minor groove of the nucleosomal DNA at SHL-3 and is angled at approximately 70° against the plane parallel to the nucleosomal disc (Figures 1B-1C). The distance between AAG and the closest histone is beyond 17 Å, and no interactions between AAG and octameric histone core were observed (Figure 1B). In addition to the protruding β-hairpin of AAG, which is inserted into the minor groove of nucleosomal DNA for direct damage recognition (Figure 1D), a few other residues also contribute to the stabilization of AAG on the nucleosome. On the damaged strand, R197, S219 and the main chain of K220 display a direct interaction with the DNA backbone (Figures 1E and 1G). On the undamaged strand, the attachment of AAG to NCP is strengthened by the interactions between the phosphate backbone and three successive positively charged residues R141, R145 and K229 (Figures 1F and 1G).

### Interaction between AAG and the AP site in the AAG-NCP^-30AP^ complex

In the AAG-NCP^-30AP^ structure, AAG shadows about 8 base-pairs of nucleosomal DNA near SHL-3 (Figure 1G). The inserted β-hairpin loop is composed of IIYGMY (residue 160-165), which makes extensive contacts with the skewed DNA and serves as the damage-recognition motif (Figures 1D and 2A and Figure S7A). The residues I160 and I161 establish hydrophobic interactions with both sides of DNA in the minor groove (Figure 2B). The aromatic residue Y162 stacks with the bases of −31A and −29T on the damaged strand through π-π interactions. Bridging residue G163 bents the terminal loop and links the β3 and β4 of AAG (Figure 2B). The main chain of G163 also contributes to the stabilization of the β-hairpin by interacting with 31T on the undamaged strand. Scaffolding residues M164 and Y165 locate in the center of the minor groove and widen the distance between two DNA backbones (Figure 2C).

**Figure 2.**
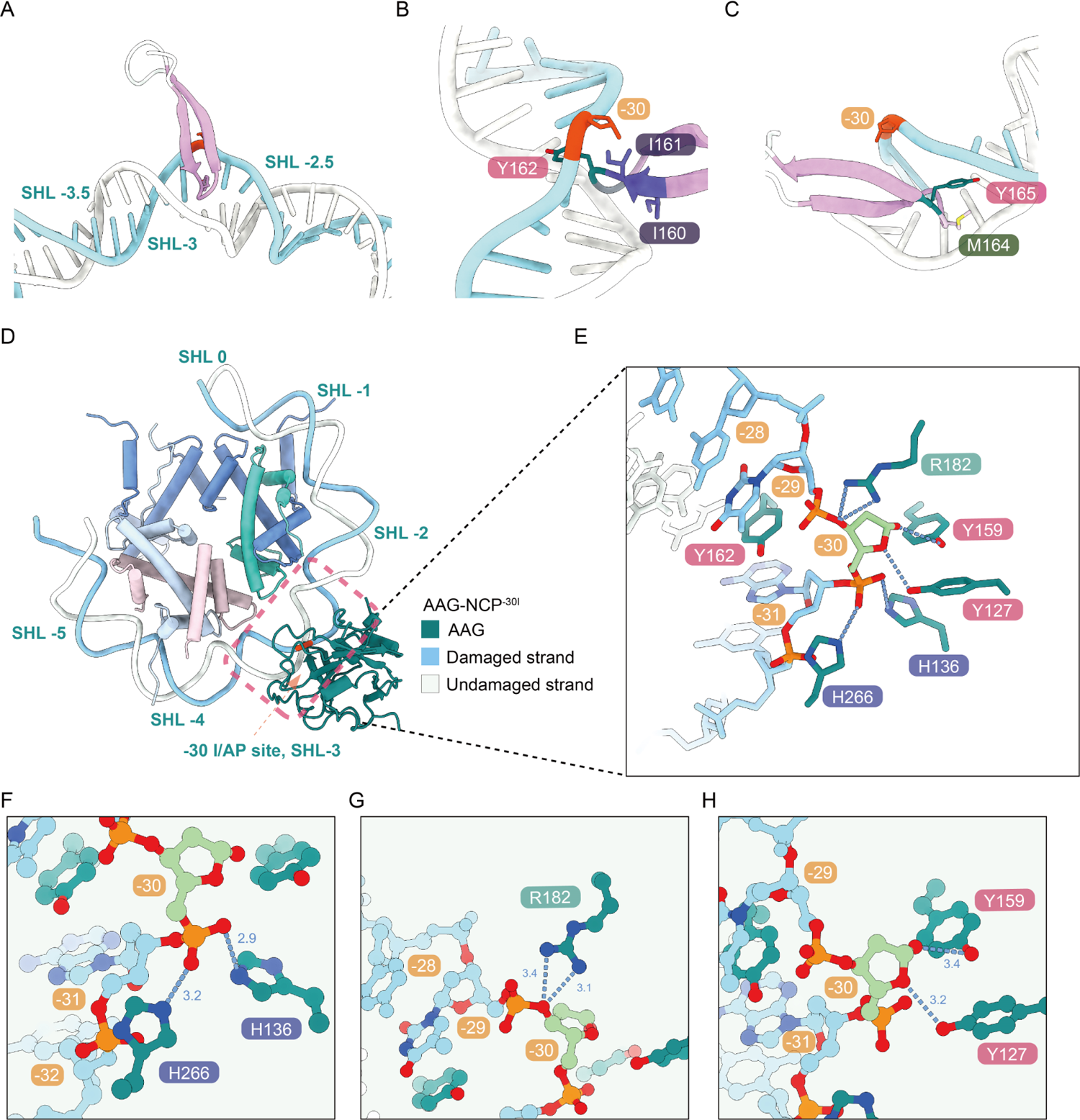
Mechanism of DNA base damage recognition by AAG in the nucleosome. **A-C**, AAG identifies the damaged base by inserting the β3-β4 hairpin into the damage located minor groove, and displaces the damaged base using Y162. **D-E**, Overview of interactions between flipped post-catalytic AP site and AAG residues (Y127, H136, Y159, R182, H266). **F-H**, Close-up views of the detailed interactions between the post-catalytic AP site and AAG.

The flipped AP site is stabilized through three sets of interactions (Figures 1G, 2D and 2E). The first interacting site consists of two basic amino acids H136 and H266 that coordinate the phosphate backbone of the AP site (Figures 2E and 2F). These two histidine residues interact with phosphate group between −30 and −31 to keep it in a proper conformation. The second site involves a positively charged residue R182 that forms a hydrogen bond with the O3’ atom of the AP site (Figures 2E and 2G). The third site involves a direct interaction of the AP-site ribose ring with two catalytically important residues, Y127 and Y159. Structural comparison of AAG-NCP^-30AP^ structure with the crystal structure of AAG bound to pyrrolidine-containing free duplex DNA (PDB: 1BNK)(*36*) revealed a different binding mode for these two residues. In the AAG-NCP^-30AP^ structure, two tyrosine residues Y127 and Y159 interact with the AP site through hydrogen bonds (Figures 2E and 2H), whereas the interaction between Y159 and the AP site is absent in the previous crystal structure, likely due to the lack of a hydroxyl group in pyrrolidine. In the crystal structure of AAG-DNA complex containing a 1,*N*^6^-ethenoadenine in the pre-catalysis state (PDB: 1EWN)(*44*), Y127 and Y159 sandwich the flipped base, and Y127 directly stacks with the modified base. Notably, most of these AP site interacting residues are invariant through evolution and a previous site-directed random mutagenesis study showed that Y127 could not be substituted and Y159 was among the lowest mutability group(*45*). Overall, AAG uses a similar set of conserved residues to interact with the DNA substrate in both the linear and nucleosomal form.

### Distortion of nucleosomal DNA in the NCP^-30I^ and AAG-NCP^-30AP^ structures

Importantly, the structures of the NCP^-30I^ and AAG-NCP^-30AP^ demonstrate apparent differences in conformation of nucleosomal DNA when compared with the structure of canonical NCP (PDB: 7OHC) (*46*). In the NCP^-30I^, the hypoxanthine base of DI pairs to pyrimidine through weaker hydrogen bonds compared to Watson-Crick base pairs, which destabilizes local DNA and leads to a global perturbation of nucleosomal DNA (Figures 3A and 3D-3E). The DI induced global perturbation, as reflected in the calculated RMSD between the damaged-strand DNA backbone of Apo NCP^-30I^ and that of the canonical NCP, shows a smallest deformation around dyad axis and increasing displacements toward nucleosomal DNA ends (Figures 3D-3E). The presence of DI at −30 does not render displacement around −30 to be largest, and the largest difference is observed at the DNA end (Figure 3E). Locally, this perturbation is characterized by an outward movement of the DNA from the histone core: The calculated RMSD of the DNA backbone from −25 to −35 (−30 +/- 5 positions) is 1.0 Å for both the damaged and intact strands (Figure 3A).

**Figure 3.**
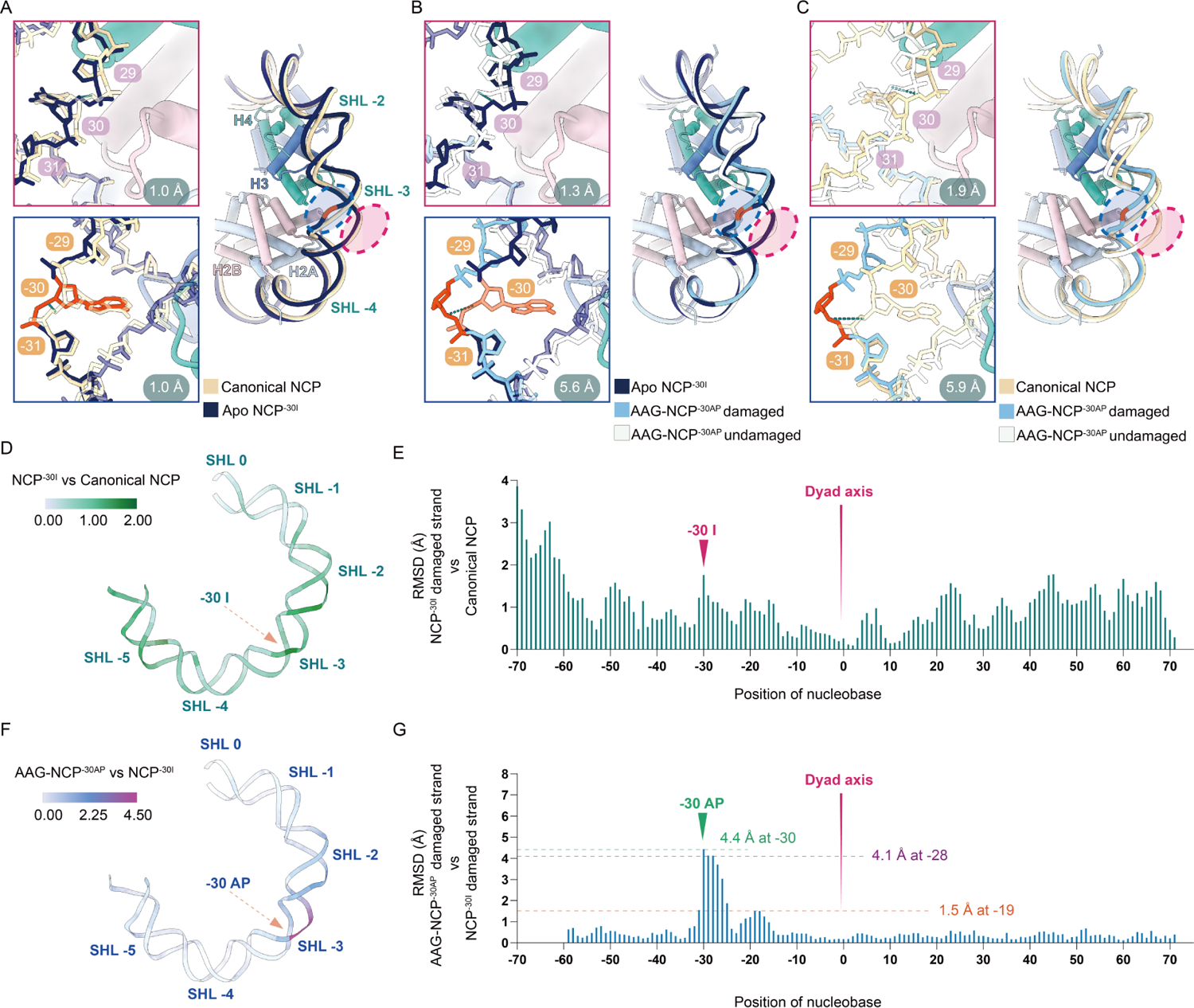
DNA distortion in NCP^-30I^ and AAG-NCP^-30AP^ structures. **A**, Nucleosomal DNA deformation in the NCP^-30I^ complex in comparison with a canonical NCP (PDB: 7OHC). Undamaged strand is labeled with red circle (right panel) and rectangle (upper left panel), and RMSD of DNA backbone from 25 to 35 is 1.0 Å. Damaged strand is labeled with blue circle (right panel) and rectangle (lower left panel), and RMSD of DNA backhone from −35 to −25 is 1.0 Å. **B**, Local DNA distortion of the AAG-NCP^-30AP^ complex in comparison with NCP^-30I^. RMSDs are 1.3 Å and 5.6 Å for undamaged and damaged strands, respectively. **C**, The collective nucleosomal DNA deformation of the AAG-NCP^-30AP^ in comparison with a canonical NCP. RMSDs are 1.9 Å and 5.9 Å for undamaged and damaged strands, reepectively. **D-E**, Temperature map and RMSD-residue plot representing deoxyinosine caused perturbation of the nucleosomal DNA calculated by comparing the NCP^-30I^ to the canonical NCP. **F-G,** Temperature map and RMSD-residue plot representing the AAG imposed local DNA distortion calculated by comparing the AAG-NCP^-30AP^ to the NCP^-30I^. Note that the terminal DNA (from −60 to exit) of the AAG-NCP^-30AP^ is relatively flexible in the map and not modelled, and therefore panel (g) only includes information from −59 to 72.

In the AAG-NCP^-30AP^, the engagement of AAG further distorts the nucleosomal DNA. Previous studies on free DNA substrate indicated that glycosylases require DNA bending at certain angle to ensure base-flipping and optimal catalysis(*47*). Comparing with the NCP^-30I^ structure, additional bending and twisting of the nucleosomal DNA around the damage site is observed, which might lead to optimal DNA bending angle for AAG engagement (Figure 3B and 3F-3G). In general, binding of AAG leads to further outward displacement of the DNA around SHL-3, and this distortion is local compared to the global perturbation caused by DI in the NCP^-30I^ (Figure 3F). The distortion out of SHL-3 rapidly decreases, and the maximum is observed at −30 with RMSD of 4.4 Å (Figure 3G). In addition, a second distortion peak was seen to be centered around −20 (SHL-2) position (Figure 3G). The calculated RMSDs around −30 AP site from −35 to −25 (−30 +/- 5 positions) are 5.6 Å and 1.3 Å for the damaged and undamaged strands, respectively (Figure 3B). A direct comparison of AAG-NCP^-30AP^ structure to the canonical NCP confirmed a very dramatic displacement of the damaged strand near SHL-3 (RMSD 5.9 Å) (Figure 3C).

Nucleosome imposes restriction to DNA through interactions between nucleosomal DNA and histones(*48*). Histone core makes contacts with DNA backbone when the minor groove of nucleosomal DNA faces histone core. Therefore, we analyzed the changes of the interactions between the histone core and the DNA backbone. The presence of DI in NCP^-30I^ causes a reduction of the buried surface area from 7141 Å^2^ to 6195 Å^2^, indicating that DI alone has significantly destabilized the interaction between nucleosomal DNA and octameric histone core (Table. 2). Further engagement of AAG has marginal effect on the overall buried surface area (6166 Å^2^), since the dramatic DNA distortion is limited to local region around damaged site (Figure 3B).

These results suggest that the damaged base induced global perturbation of nucleosomal DNA destabilizes the nucleosome and relives nucleosome-imposed restriction, which likely plays a determinant role in AAG recruitment. Due to the high solution accessibility of −30, additional local DNA distortion induced by AAG alone is sufficient to deform nucleosomal DNA and allow AAG engagement to the damaged site.

### Distortion of nucleosomal DNA in the NCP^-50I^ and AAG-NCP^-50AP^ structures

We next examined nucleosomes with a damaged base at different SHL (SHL-5) to study the effect of translational position. In the AAG-NCP^-50AP^ complex, a DI was introduced into the position of −50 at SHL-5, which could be regarded as a roughly equivalent position of −30 in terms of solution accessibility (Figures 4A-4B). Subsequently, the structures of Apo NCP^-50I^ and AAG-NCP^-50AP^ complexes were solved at resolutions of 2.9 Å and 3.0 Å, respectively (Figures S3, S7 and S8A-S8B).

**Figure 4.**
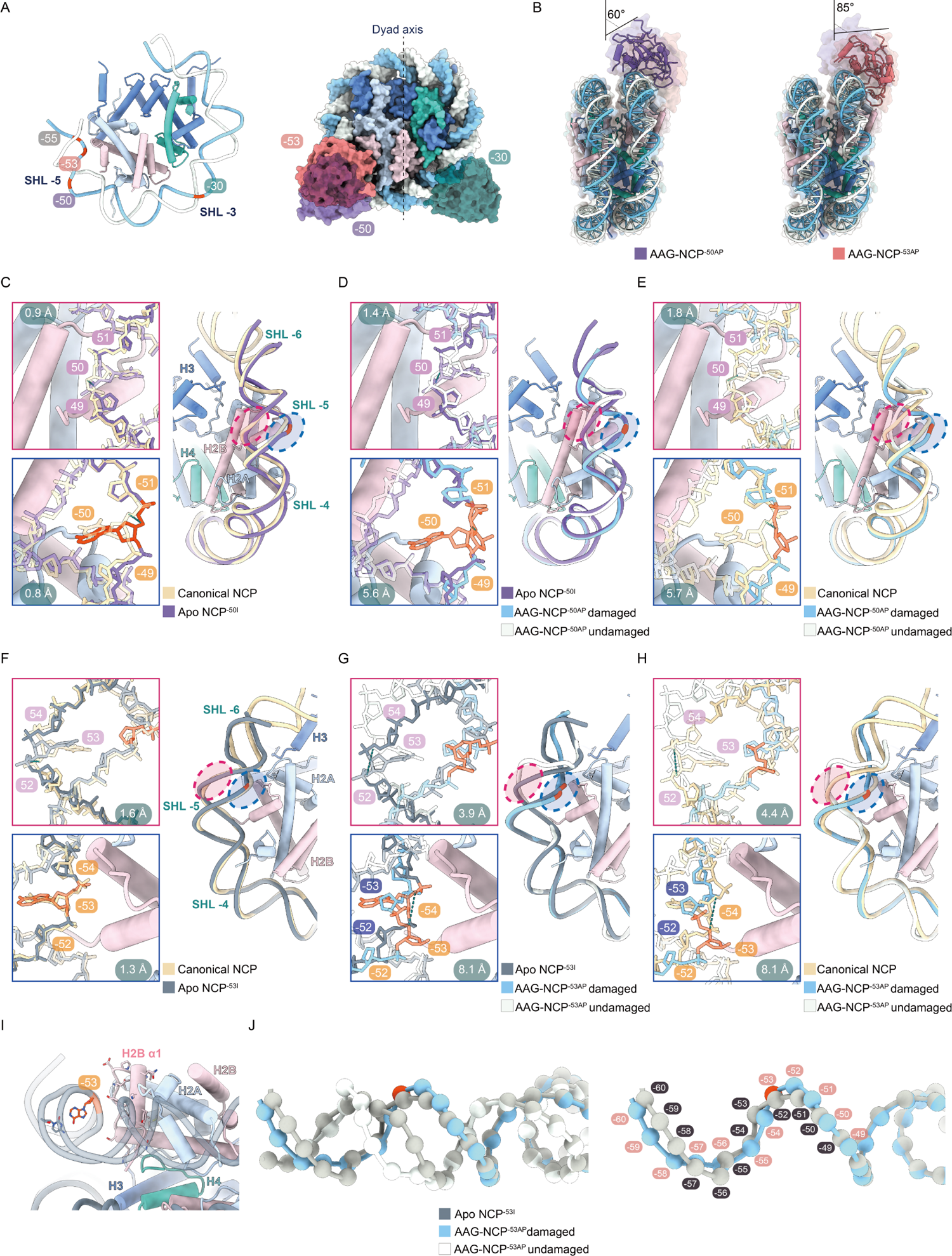
Structural comparison between NCP^-50I^, AAG-NCP^-50AP^, NCP^-53I^, AAG-NCP^-53AP^ and the canonical NCP. **A**, Superimposition of the models of the AAG-NCP^-30AP^, AAG-NCP^-50AP^ and AAG-NCP^-53AP^ to highlight the orientation difference of AAG the three structures. **B**, Comparison of the AAG-NCP^-50AP^ and AAG-NCP^-53AP^. **C**, The nucleosomal DNA perturbation in the NCP^-50I^ in comparison with a canonical NCP (PDB: 7OHC). RMSD of the undamaged DNA backbone from 45 to 55 is 0.9 Å, and 0.8 Å for the undamaged strand from −55 to −45. **D,** Local DNA distortion of the AAG-NCP^-50AP^ in comparison with the NCP^-50I^. RMSDs are 1.4 Å and 5.6 Å for the undamaged and damaged strands, respectively. **E,** The nucleosomal DNA deformation of the AAG-NCP^-50AP^ in comparison with a canonical NCP. RMSDs are 1.8 Å and 5.7 Å for the undamaged and damaged strands, respectively **F.** The nucleosomal DNA perturbation in the NCP^-53I^ in comparison with a canonical NCP (PDB: 7OHC). RMSD of the undamaged DNA backbone from 48 to 58 is 1.6 Å, and 1.3 Å for the undamaged strand from −58 to −48. **G,** In the AAG AAG-NCP^-53AP^, drastic local DNA distortion is accompanied by local register shift. In comparison with the NCP^-53I^, RMSD of the undamaged strand is 3.9 Å, and the RMSD of the damaged strand is 8.1 Å. AP site at −53 is relocated to −52 in relative to the register of the NCP^-53I^. **H,** The nucleosomal DNA deformation in the AAG-NCP^-53AP^ in comparison with a canonical NCP^-53I^. RMSDs are 4.4 Å and 8.1 Å for the undamaged and damaged strands, respectively. **I,** The original DNA register of the NCP^-53I^ prohibits base flipping at −53, due to steric clashe with histone H2B. **J,** The local register shift of the AAG-NCP^-53AP^ in comparison to the register of the NCP^-53I^.

Comparison of the Apo NCP^-50I^ to a canonical NCP (PDB: 7OHC) (*46*) reveals a similar pattern of global perturbation as seen in the structure of the NCP^-30I^ (Figure S9A). The presence of DI at −50 causes a noticeable movement of both strands away from the histone core: at −55 to −45 (−50 +/- 5 positions) of SHL-5, the calculated RMSDs for the DNA backbone are 0.8 Å and 0.9 Å for the damaged and undamaged strands, respectively (Figure 4C).

In the AAG-NCP^-50AP^ structure, the engagement of AAG is similar to that in the AAG-NCP^-30AP^ in general, although AAG in AAG-NCP^-50AP^ is now angled at approximately 60° against the plane parallel to the nucleosome disc (Figure 4B). Globally, the buried surface area between the DNA and the histone core decreases from 7134 Å^2^ to 6248 Å^2^ in the presence of DI at −50, and further decrease to 5977 Å^2^ after engaging with AAG (Table. 2). Further DNA distortion was also observed upon AAG engagement, and displays a predominantly local effect: The maximal distortion is observed at −50 with RMSD of 3.0 Å (Figure S9B), and the average displacements for the damaged and undamaged strands at −55 to −45 of SHL-5 (−50 +/-5 positions) are 5.6 Å and 1.4 Å, respectively (Figure 4D). When a canonical NCP was used for comparison, the DNA backbone RMSDs at SHL-5 are 5.7 Å and 1.8 Å for the damaged and undamaged strands, respectively (Figure 4E).

This degree of local DNA distortion in the AAG-NCP^-50AP^ is less than that in the AAG-NCP^-30AP^, but the overall similarity suggests a general mechanism for AAG to engage with DNA base damage in solvent-facing positions, which leverages on the global structural perturbation of the nucleosome by DI nucleotide.

### Structure of the AAG-NCP**^-53AP^** complex containing an occluded damaged base with medium solution accessibility

Within a SHL region of the nucleosomal DNA, damaged bases in different rotational orientations should have sharply different solution accessibility, and thus different accessibility for repair proteins. Although the DI in the NCP^-53I^ is at an occluded position of SHL-5 with medium solution accessibility, (Figures 1A and 4A), our data showed that the assembly of the AAG-NCP^-53^ complex displays a comparable high efficiency as the −30 and −50 positions (Figure S4A). We subsequently determined the structures of the NCP^-53I^ at 2.8 Å and AAG-NCP^-53AP^ at 3.1 Å (Figures S4, S7 and S8C-S8D). Comparison of the NCP^-53I^ to a canonical NCP (PDB: 7OHC) reveals a similar global perturbation as in the NCP^-30I^ and NCP^-50I^ (Figure S9C), featuring a local outward movement of the nucleosomal DNA around the damage site, with the calculated RMSDs of the DNA backbone for the damaged and undamaged strands (−53 +/- 5 positions) being 1.3 Å and 1.6 Å, respectively (Figure 4F).

AAG in AAG-NCP^-53AP^ is angled at approximately 85° against the plane parallel to the nucleosome disc (Figure 4B). Importantly, in the AAG-NCP^-53AP^, a drastic local DNA distortion was observed (Figures 4G and 4H), and the maximal distortion is at −53 with RMSD of 7.4 Å (Figure S9D). There is a significant widening of the minor groove in the local region of −53 position compared to the NCP^-53I^ (Figure 4J), which is also reflected by much larger displacements for both the damaged and undamaged strands: The calculated RMSDs of the DNA backbones from −58 to −48 (−53 +/- 5 positions) are 8.1 Å and 3.9 Å for the damaged and undamaged strands, respectively (Figure 4G). When comparing the AAG-NCP^-53AP^ with a canonical NCP (PDB: 7OHC), the RMSDs for the two strands are 8.1 Å and 4.4 Å, respectively (Figure 4H). Globally, the buried surface area between the DNA and the histone core decreases from 7141 Å^2^ to 6075 Å^2^ in the presence of DI at −53, and further decrease to 5897 Å^2^ after engaging with AAG (Table. 2).

Very intriguingly, in comparison to the local DNA distortion in the AAG-NCP^-30AP^ and AAG-NCP^-50AP^ (Movie. 1, 2) an additional translocation of local DNA around −53 was observed (Movie. 3, 4). The AP site at −53 in the AAG-NCP^-53AP^ is now at −52 in spatial relation to the translational register of NCP^-53I^. Since base-flipping at −53 on a regular NCP would cause clash with H2B, this local register shift from −53 to −52 appears to be necessary for AAG to interact with flipped base (Figure 4I). It must be noted that the global translational register is not affected, and this translocation is restricted to the region from the damage site to the near exit end of the nucleosomal DNA.

Therefore, these observations suggest that for positions with medium solution accessibility, AAG could induce drastic local DNA distortion that allows local register shift to increase the solution accessibility of occluded damaged base (Figure 4J). This structural finding explains a previous *in vitro* data that although −53 is an occluded position with medium solution accessibility, AAG is highly active at this position(*31*).

### Partial opening of the nucleosome in the AAG-NCP^-55^ complex

Different from the previous DI-containing NCPs, the DI in the AAG-NCP^-55AP^ is in a completely buried position with low solution accessibility. Therefore, one would expect a very low activity of AAG at this site. To our surprise, the NCP^-55I^ is still capable of forming a stable complex with AAG, as shown in the glycerol density gradient experiment (Figure S5). Similarly, cryo-EM analysis revealed two major conformational populations for the AAG-NCP^-55^ sample. One is the structure of NCP^-55I^ determined at 2.8 Å (Figures S5, S7 and S8E), and the other one is an unusual NCP structure at 2.9 Å resolution, with the terminal nucleosomal DNA highly flexible, unresolved in the map (From SHL-5 to the proximal end) (Figure 5A). In this unusual structure, however, density of AAG could not be found. Given the strong association of AAG with the NCPs in our biochemical preparation (Figure S5), it is highly likely this structure in fact reflects the AAG-NCP^-55AP^ complex, in which a drastic conformational change of the DNA at the region of −55 has caused the opening of the terminal DNA duplex (Figure 5B). This assignment is also supported by previous data that AAG still retained ∼50% activity towards a damaged base at −55 position(*31*).

**Figure 5.**
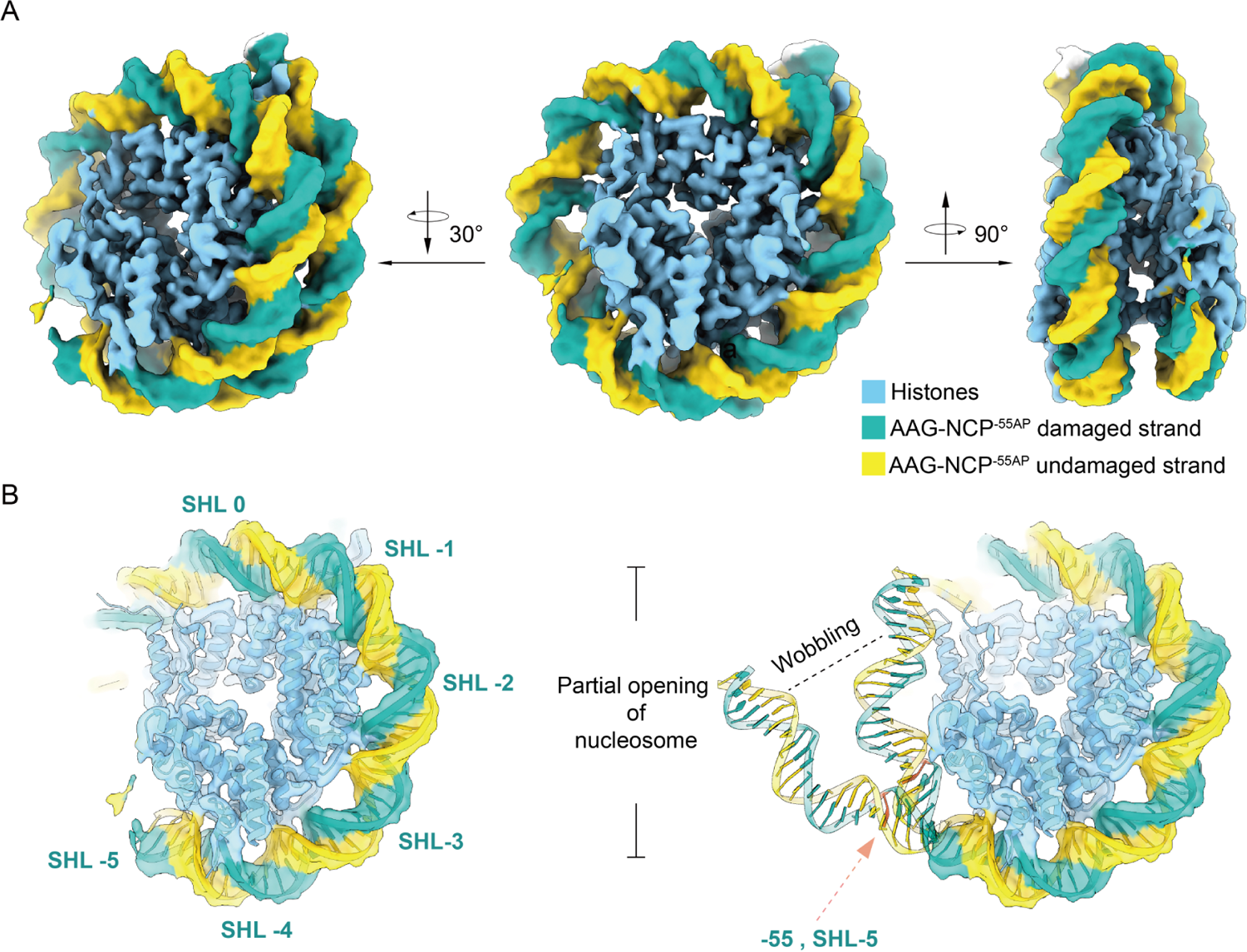
Partial opening of nucleosome in the AAG-NCP^-55AP^. **A**, Cryo-EM map of the AAG-NCP^-55AP^ complex. **B,** Partial opening of nucleosome in the AAG-NCP-55AP. DNA from SHL-5 to proximal end is peeled off from the histone core.

Similarly, a global perturbation of nucleosomal DNA by DI in NCP^-55I^ was observed, but the proximal end of the DNA displays a much larger displacement (Figure S9E). The buried surface area between the DNA and the histone core decreases from 7517 Å^2^ to 6745 Å^2^ in the presence of DI at −55 (Table.2). Locally at −60 to −50 (55 +/- 5 positions), the −55 DI induced structural displacements of DNA backbone are 1.1 Å and 1.3 Å for the damaged and undamaged strands, respectively, which is in a similar range as the NCP^-30I^, NCP^-50I^ and NCP^-53I^ complexes (Figure S10).

It is of very low possibility that this unusual NCP is still in the pre-catalytic state before AAG binding. The dynamic nature of nucleosome could lead to spontaneous unwrapping of nucleosomal DNA(*32–34*), and the significant destabilization of the DNA at the exit observed in the NCP^-55I^ could further increases the unwrapping possibility. In this scenario, the partial opening of nucleosomal DNA gives a short-lived window for AAG to access the damaged site. The binding of AAG to the detached DNA duplex renders this process less reversible, allowing the formation of a stable AAG-NCP^-55^ complex for subsequent excision reaction.

Altogether, the presence of a DI in a completely buried position also leads to a global perturbation of the nucleosomal DNA. When the damage base is near the exit, the perturbation could increase the possibility of terminal DNA duplex opening to facilitate AAG engagement.

## Discussion

Over the past several decades, immense amount of effort has been devoted to the elucidation of principles governing interactions between DNA-binding proteins and free DNA duplex (reviewed in (*49, 50*)). However, in a more physiological context, how DNA-binding proteins interrogate nucleosomes to detect and read hidden information on chromatin is still not completely clear(*51*). Recent structural studies on nucleotide excision repair (*52*), retroviral integration(*53, 54*) and pioneer transcription factors(*55, 56*) have revealed that these factors could employ a few different mechanisms, such as nucleosomal DNA register shifting and local DNA distortion, to induce nucleosome deformation to fulfill their molecular functions.

In the present work, we determined a set of cryo-EM structures of the NCPs and AAG-NCP complexes with a damage mimicking DI nucleotide placed in different representative positions on nucleosome. Although AAG in these complexes are positioned differently on the nucleosomes, the binding mode of AAG on the nucleosome is generally similar to AAG on free DNA duplex(*36, 44*), with only limited interactions through several evolutionarily invariant residues with the negatively charged DNA backbone(*45*) (Figure 1G). These structures also show a highly conserved mechanism for AAG to recognize the damaged base, involving the insertion of a protruding β hairpin into the minor groove of the damage site, the replacement of the damaged base by a tyrosine residue, and stabilization of the flipped base/AP site with two aromatic residues (Figures 1D and 2). These similarities suggest that a crucial and potentially rate-limiting step for base excision reactions could be the recruitment of AAG onto the damage sites.

Interestingly, with pair-wise comparison to the canonical NCP, we discovered that the presence of a single DI nucleotide alone is sufficient to perturb nucleosomal DNA globally (Figures 3D-3E and Figures S9A, S9C and S9E), resulting in a collectively reduced buried surface area between nucleosomal DNA and histone core, by 10-15% reduction depending on the DI position (Table, 2). This global perturbation of nucleosomal DNA is generally in a similar pattern regardless of the position of the damaged base (Figures 3D-3E, Figures S9A, S9C and S9E). The DNA deformation caused by DI is most apparent near the exit of nucleosomal DNA, and the perturbation attenuates toward the center of the nucleosomal DNA and reaches its minimum at the dyad axis (Figures 3D-3E, Figures S9A, S9C and S9E).

After formation of the stable AAG-NCP^AP^ complex, the binding of AAG induces a dramatic but relatively local distortion on nucleosomal DNA around the damaged base (Figures 3F, 3G and Figures S9B, S9D). Dependent on the translational and rotational positions of the damaged site, AAG makes use of distinct mechanisms to get access to the damaged base. For the DI in solvent-facing positions with high solution accessibility, where AAG is highly active(*31*) and has a direct access to these positions, AAG directly augments the local DNA distortion to recognize the damage site (Figure S11A). For the DI in occluded positions such as −53 with medium accessibility, where AAG activity is expected to be lower comparing to solvent-facing positions(*31*), AAG induces drastic local DNA distortion, including both twisting and translocation of DNA to generate a shift of local DNA register by 1-bp to relieve the nucleosome-imposed spatial hindrance for accessing the damaged site (Figures 4I, 4J and Figure S11B). As for deeply embedded positions (for example, −55 in our study), local DNA distortion and limited register shift are insufficient to fully expose the buried base. Our results suggest that globally perturbed nucleosome by DI might be more prone to spontaneous unwrapping(*34*). AAG can make use of this feature to capture the detached terminal DNA duplex and render this process less reversible to favor the unwrapping direction. Therefore, in these completely buried sites, a partial opening of the nucleosomal DNA might be a prerequisite for the recruitment and catalysis of AAG (Figure 5B, and Figure S11C).

Altogether, in all these above-mentioned scenarios, the altered structural dynamics of the nucleosome likely plays a major role in recruiting AAG to the damages site. The global perturbation of the nucleosomal DNA by DI nucleotide simply reflects the fact that disruption of a single base pair could weaken DNA-histone interaction and alter the conformational landscape of the nucleosome. In a kinetics view, this would result in an increased sampling of the otherwise less possible conformations of the nucleosome. AAG is capable of capturing these transient conformations and forms a stable AAG-NCP complex for the subsequent excision reaction. In summary, our work reveals the effect of damaged base to nucleosome stability and provide a mechanistic framework for understanding how DNA glycosylase AAG exploits the structural dynamics of nucleosome to engage with DNA base damage in the nucleosome. In a broader context, our work also contributes to the general knowledge of how the structural dynamics of nucleosome inter-plays with DNA-binding proteins to regulate their actions on chromatinized eukaryotic genome.

## Supplementary Materials

### Methods

#### Construct design and protein expression

Gene encoding the N-terminus truncated sequence of human AAG (residue 80-298) was cloned into pET22b vector with an N-terminal 6× His tag. Plasmid verified by sequencing was transformed into BL21 (DE3) *E. coli* cells for overexpression. At OD_600_ of 0.6, expression of AAG was induced by 0.1 mM isopropyl β-D-1-thiogalactopyranoside (IPTG) at 18°C for 17 hours. Cells were collected by centrifugation (3500 rpm, 10 min, 25°C) and resuspended in buffer A (20 mM Tris-HCl, pH 7.5, 100 mM NaCl). The suspension was flash-frozen in liquid nitrogen and stored at −80 °C for purification.

#### Purification of AAG

All steps were carried out at 4 °C. Frozen cells were thawed and lysed by sonication in buffer A supplemented with 0.1 mM phenylmethanesulfonylfluoride (PMSF). The lysate was then supplemented with 0.5 U/mL benzonase nuclease and centrifuged at 18,000 rpm for 40 min. The supernatant was filtered with a 0.45 μm syringe filter, then loaded onto 5 mL Ni-smart beads 6FF FPLC column (Smart-Lifesciences, Changzhou), and eluted with a linear gradient of imidazole using buffer A and buffer B (20 mM Tris-HCl, pH 7.5, 100 mM NaCl, 1 M Imidazole). Peak fractions were collected and pooled for dialysis in buffer C (20 mM Tris-HCl pH 7.5, 100 mM NaCl, 1 mM EDTA, 1 mM DTT) overnight to remove imidazole. Dialyzed sample was centrifuged to remove any possible precipitates, and the supernatant was loaded onto 5 mL Heparin beads 6FF FPLC column (Smart-Lifesciences, Changzhou) pre-equilibrated with buffer C, and then eluted with a linear gradient of buffer C and buffer D (20 mM Tris-HCl pH 7.5, 1 M NaCl, 1 mM EDTA, 1 mM DTT). Peak fractions were pooled and concentrated, loaded onto Superdex 75 10/300 GL (Cytiva) column pre-equilibrated in buffer C. Peak fractions were collected and analyzed by SDS-PAGE. The AAG proteins were concentrated to 1 mg/mL and stored at −80 °C in small aliquots.

#### Widom 601 DNA preparation

Widom 601 top strand sequence: CCTGGAGAATCCCGGTGCCGAGGCCGCTCAATTGGTCGTAGACAGCTCTAGCACCGCTTAAACGCACGTACGCGCTGTCCCCCGCGTTTTAACCGCCAAGGGGATTACTCCCTAGTCTCCAGGCACGTGTCAGATATATACATCCTGTGCA T

Widom 601 −30I bottom strand sequence: ATGCACAGGATGTATATATCTGACACGTGCCTGGAGACTAGGGAGTA**I**TC CCCTTGGCGGTTAAAACGCGGGGGACAGCGCGTACGTGCGTTTAAGCGGT GCTAGAGCTGTCTACGACCAATTGAGCGGCCTCGGCACCGGGATTCTCCA GG

Widom 601 −50I bottom strand sequence: ATGCACAGGATGTATATATCTGACACG**I**GCCTGGAGACTAGGGAGTAATC CCCTTGGCGGTTAAAACGCGGGGGACAGCGCGTACGTGCGTTTAAGCGGT GCTAGAGCTGTCTACGACCAATTGAGCGGCCTCGGCACCGGGATTCTCCA GG

Widom 601 −53I bottom strand sequence: ATGCACAGGATGTATATATCTGAC**I**CGTGCCTGGAGACTAGGGAGTAATC CCCTTGGCGGTTAAAACGCGGGGGACAGCGCGTACGTGCGTTTAAGCGGT GCTAGAGCTGTCTACGACCAATTGAGCGGCCTCGGCACCGGGATTCTCCA GG

Widom 601 −55I bottom strand sequence: ATGCACAGGATGTATATATCTG**I**CACGTGCCTGGAGACTAGGGAGTAATC CCCTTGGCGGTTAAAACGCGGGGGACAGCGCGTACGTGCGTTTAAGCGGT GCTAGAGCTGTCTACGACCAATTGAGCGGCCTCGGCACCGGGATTCTCCA GG

The widom 601 sequence(*57*) was inserted into pET-51b vector, and the resulting plasmid was used as PCR template. The plasmid was transformed into Trans1-T1 *E. coli* cells and isolated using Plasmid Mini Kit I (Omega Bio-Tek). Damaged base containing Widom 601 DNA was prepared by PCR with deoxyinosine containing primers. The primers used in this study were listed in (Table 1). A typical purification procedure required 10 mL of PCR reaction product. PCR reaction product was loaded onto 5 mL Q beads 6FF FPLC column (Smart-Lifesciences, Changzhou) pre-equilibrated in 20 mM Tris-HCl (pH 8.0), eluted with linear gradient of NaCl (from 0 to 2 M, 20 mM Tris-HCl pH 8.0). Peak fractions containing Widom 601 DNA were collected and concentrated using 10K AmiconUltra-15 centrifugal filter unit (Merck).

**Table 1.**
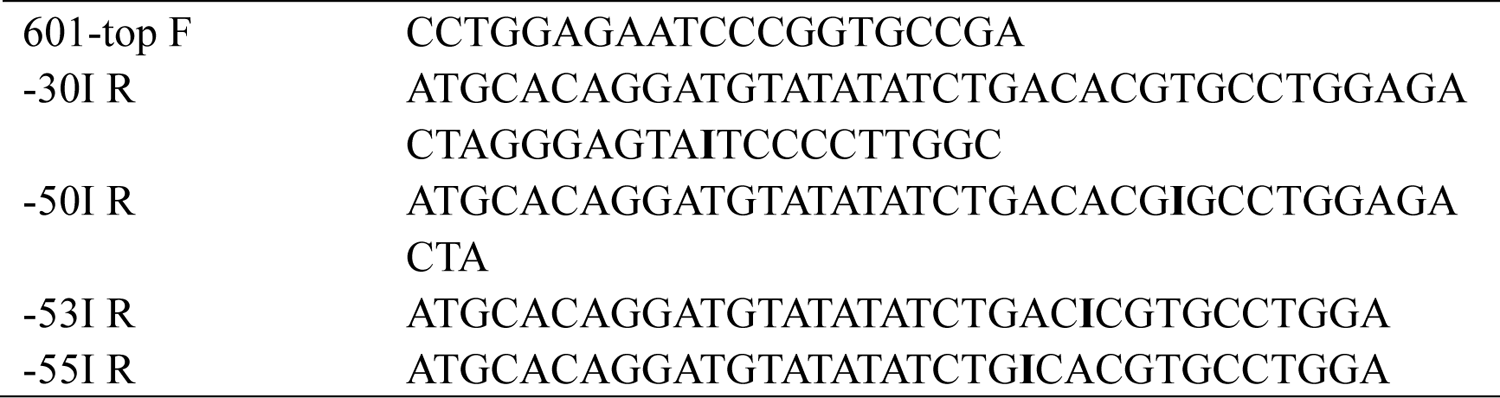
Sequences of the oligonucleotides used as primers for PCR amplification

**Table 2.**
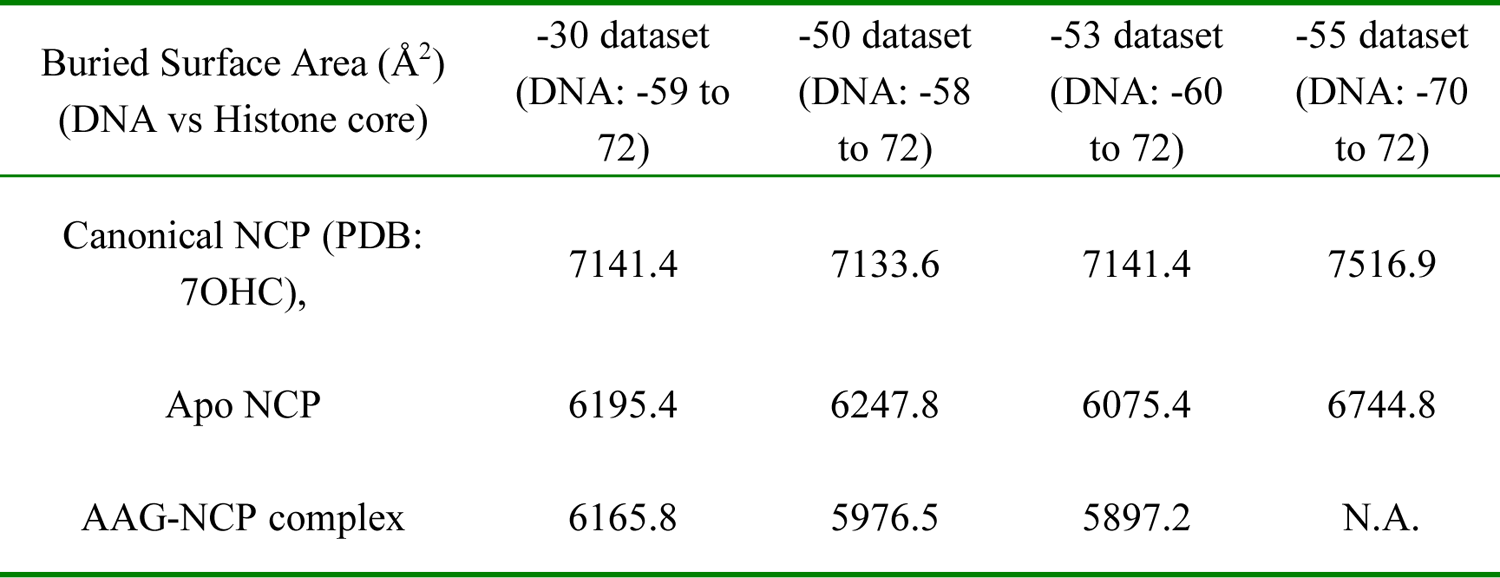
Buried surface area between nucleosomal DNA and Octameric histone core.

**Table 3.**
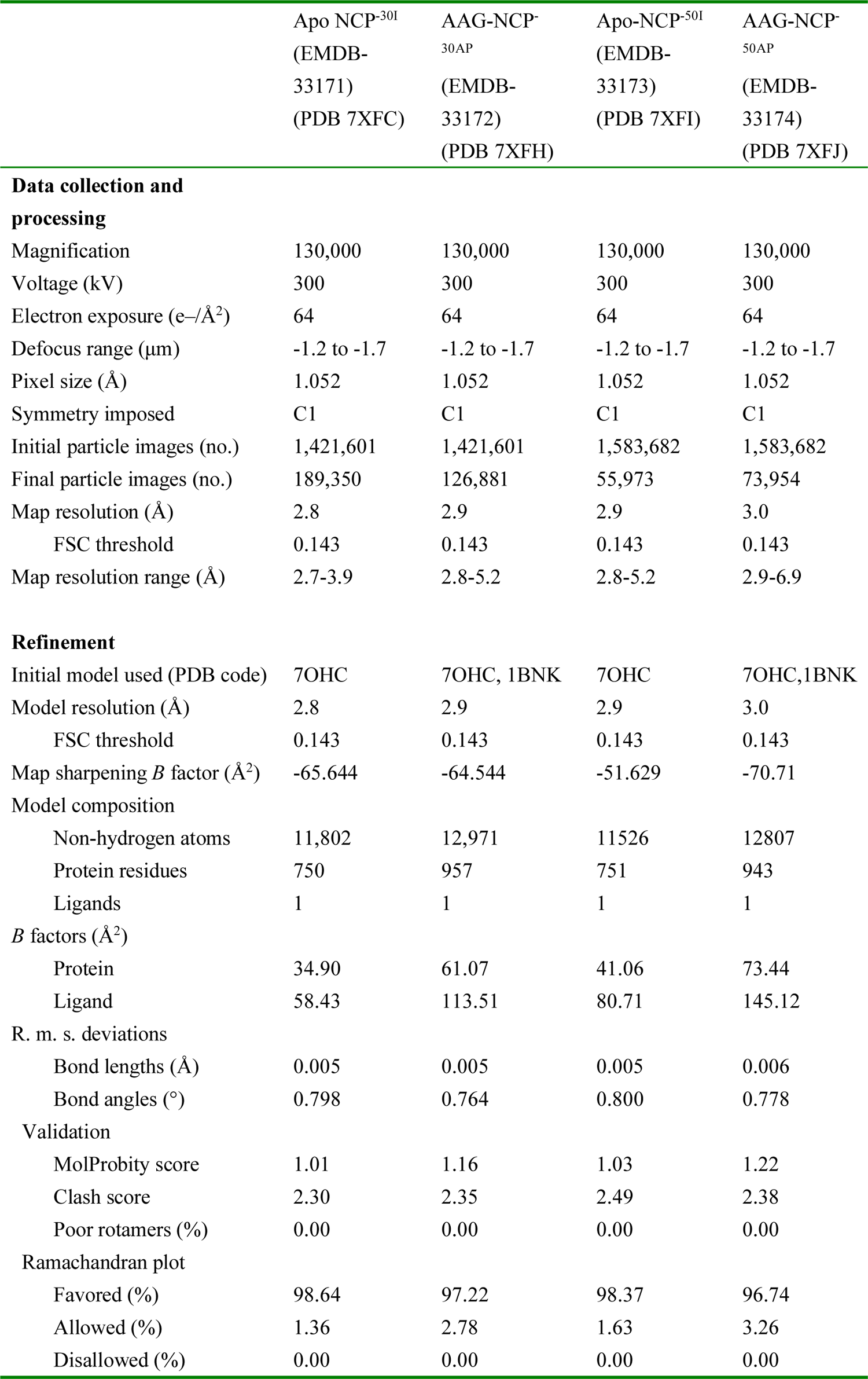
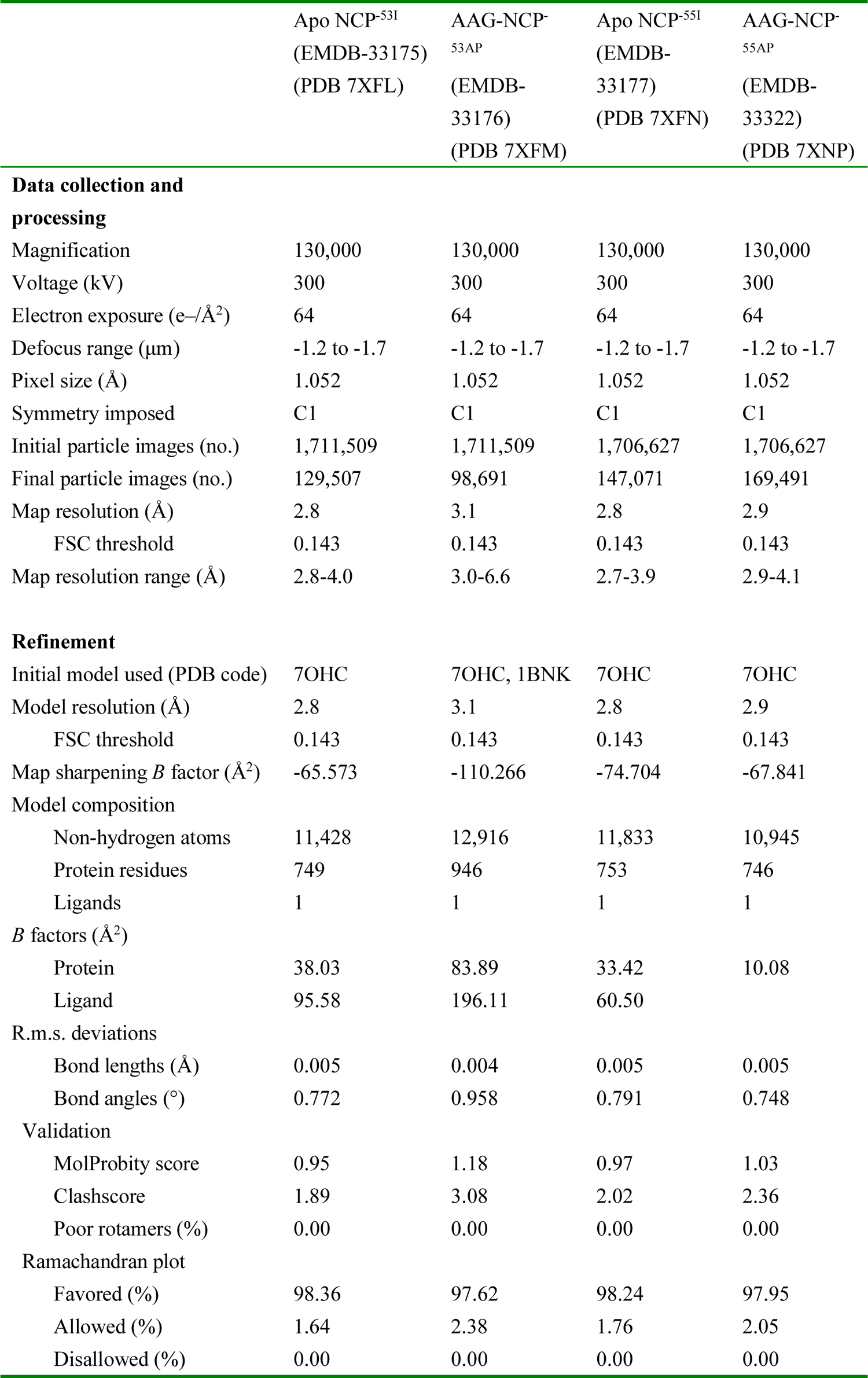
Cryo-EM data collection, refinement and validation statistics

|  | Apo NCP<sup>-30I</sup><br>(EMDB-33171)<br>(PDB 7XFC) | AAG-NCP<sup>-30AP</sup><br>(EMDB-33172)<br>(PDB 7XFH) | Apo-NCP<sup>-50I</sup><br>(EMDB-33173)<br>(PDB 7XFI) | AAG-NCP<sup>-50AP</sup><br>(EMDB-33174)<br>(PDB 7XFJ) |
| --- | --- | --- | --- | --- |
| **Data collection and processing** |  |  |  |  |
| Magnification | 130,000 | 130,000 | 130,000 | 130,000 |
| Voltage (kV) | 300 | 300 | 300 | 300 |
| Electron exposure (e-/Å<sup>2</sup>) | 64 | 64 | 64 | 64 |
| Defocus range (μm) | -1.2 to -1.7 | -1.2 to -1.7 | -1.2 to -1.7 | -1.2 to -1.7 |
| Pixel size (Å) | 1.052 | 1.052 | 1.052 | 1.052 |
| Symmetry imposed | C1 | C1 | C1 | C1 |
| Initial particle images (no.) | 1,421,601 | 1,421,601 | 1,583,682 | 1,583,682 |
| Final particle images (no.) | 189,350 | 126,881 | 55,973 | 73,954 |
| Map resolution (Å) | 2.8 | 2.9 | 2.9 | 3.0 |
| FSC threshold | 0.143 | 0.143 | 0.143 | 0.143 |
| Map resolution range (Å) | 2.7-3.9 | 2.8-5.2 | 2.8-5.2 | 2.9-6.9 |
| **Refinement** |  |  |  |  |
| Initial model used (PDB code) | 7OHC | 7OHC, 1BNK | 7OHC | 7OHC, 1BNK |
| Model resolution (Å) | 2.8 | 2.9 | 2.9 | 3.0 |
| FSC threshold | 0.143 | 0.143 | 0.143 | 0.143 |
| Map sharpening *B* factor (Å<sup>2</sup>) | -65.644 | -64.544 | -51.629 | -70.71 |
| Model composition |  |  |  |  |
| Non-hydrogen atoms | 11,802 | 12,971 | 11526 | 12807 |
| Protein residues | 750 | 957 | 751 | 943 |
| Ligands | 1 | 1 | 1 | 1 |
| *B* factors (Å<sup>2</sup>) |  |  |  |  |
| Protein | 34.90 | 61.07 | 41.06 | 73.44 |
| Ligand | 58.43 | 113.51 | 80.71 | 145.12 |
| R. m. s. deviations |  |  |  |  |
| Bond lengths (Å) | 0.005 | 0.005 | 0.005 | 0.006 |
| Bond angles (°) | 0.798 | 0.764 | 0.800 | 0.778 |
| Validation |  |  |  |  |
| MolProbity score | 1.01 | 1.16 | 1.03 | 1.22 |
| Clash score | 2.30 | 2.35 | 2.49 | 2.38 |
| Poor rotamers (%) | 0.00 | 0.00 | 0.00 | 0.00 |
| Ramachandran plot |  |  |  |  |
| Favored (%) | 98.64 | 97.22 | 98.37 | 96.74 |
| Allowed (%) | 1.36 | 2.78 | 1.63 | 3.26 |
| Disallowed (%) | 0.00 | 0.00 | 0.00 | 0.00 |

|  | Apo NCP<sup>-53I</sup><br>(EMDB-33175)<br>(PDB 7XFL) | AAG-NCP<sup>-53AP</sup><br>(EMDB-33176)<br>(PDB 7XFM) | Apo NCP<sup>-55I</sup><br>(EMDB-33177)<br>(PDB 7XFN) | AAG-NCP<sup>-55AP</sup><br>(EMDB-33322)<br>(PDB 7XNP) |
| --- | --- | --- | --- | --- |
| **Data collection and processing** |  |  |  |  |
| Magnification | 130,000 | 130,000 | 130,000 | 130,000 |
| Voltage (kV) | 300 | 300 | 300 | 300 |
| Electron exposure (e-/Å<sup>2</sup>) | 64 | 64 | 64 | 64 |
| Defocus range (μm) | -1.2 to -1.7 | -1.2 to -1.7 | -1.2 to -1.7 | -1.2 to -1.7 |
| Pixel size (Å) | 1.052 | 1.052 | 1.052 | 1.052 |
| Symmetry imposed | C1 | C1 | C1 | C1 |
| Initial particle images (no.) | 1,711,509 | 1,711,509 | 1,706,627 | 1,706,627 |
| Final particle images (no.) | 129,507 | 98,691 | 147,071 | 169,491 |
| Map resolution (Å) | 2.8 | 3.1 | 2.8 | 2.9 |
| FSC threshold | 0.143 | 0.143 | 0.143 | 0.143 |
| Map resolution range (Å) | 2.8-4.0 | 3.0-6.6 | 2.7-3.9 | 2.9-4.1 |
| **Refinement** |  |  |  |  |
| Initial model used (PDB code) | 7OHC | 7OHC, 1BNK | 7OHC | 7OHC |
| Model resolution (Å) | 2.8 | 3.1 | 2.8 | 2.9 |
| FSC threshold | 0.143 | 0.143 | 0.143 | 0.143 |
| Map sharpening *B* factor (Å<sup>2</sup>) | -65.573 | -110.266 | -74.704 | -67.841 |
| Model composition |  |  |  |  |
| Non-hydrogen atoms | 11,428 | 12,916 | 11,833 | 10,945 |
| Protein residues | 749 | 946 | 753 | 746 |
| Ligands | 1 | 1 | 1 | 1 |
| *B* factors (Å<sup>2</sup>) |  |  |  |  |
| Protein | 38.03 | 83.89 | 33.42 | 10.08 |
| Ligand | 95.58 | 196.11 | 60.50 |  |
| R.m.s. deviations |  |  |  |  |
| Bond lengths (Å) | 0.005 | 0.004 | 0.005 | 0.005 |
| Bond angles (°) | 0.772 | 0.958 | 0.791 | 0.748 |
| Validation |  |  |  |  |
| MolProbity score | 0.95 | 1.18 | 0.97 | 1.03 |
| Clashscore | 1.89 | 3.08 | 2.02 | 2.36 |
| Poor rotamers (%) | 0.00 | 0.00 | 0.00 | 0.00 |
| Ramachandran plot |  |  |  |  |
| Favored (%) | 98.36 | 97.62 | 98.24 | 97.95 |
| Allowed (%) | 1.64 | 2.38 | 1.76 | 2.05 |
| Disallowed (%) | 0.00 | 0.00 | 0.00 | 0.00 |

Concentrated solution was supplemented with 1/10 volume of 3 M sodium acetate and 2 volumes of 100% ethanol to precipitate DNA. After 1 hour of freezing at −40 °C, the mixture was centrifuged at 15000 rpm at 4 °C for 10 min, and the DNA pellet was washed with ice cold 70% ethanol and dried in air. The resulting DNA pellet was stored at −80°C for nucleosome reconstitution.

#### Histone octamer preparation

Method for expression and purification of *Xenopus laevis* histones were adapted from previous studies(*41, 43*). In brief, H2A from *Xenopus laevis* was cloned into pET-22b vector, which contained a 6× His tag and a TEV site at N-terminus of H2A. H2B, H3 and H4 were cloned into pET-3a vectors.

Histone expression was induced with IPTG at 3*7*°C, and histone expressing bacterial cells from 1L H2A, 4L H2B, 2L H3 and 2L H4 of bacterial cultures were mixed to achieve stoichiometry of 1:1:1:1. Mixed cells were lysed by sonication, and inclusion body was recovered by centrifugation and dissolved in 20 mM acetate pH 5.2, 8 M guanidine hydrochloride, 10 mM DTT. Supernatants containing denatured histones were recovered by centrifugation, and transferred into dialysis bags for histone octamer refolding. Histone octamer was dialyzed against with a buffer (20 mM Tris pH 8.0, 2 M NaCl, and 2 mM β-mercaptoethanol) at 4°C for three times, with each time more than 8 hours. After dialysis, supernatant containing refolded histone octamer was centrifugated, loaded onto 5 mL Ni-smart beads 6FF FPLC column (Smart-Lifesciences, Changzhou), and eluted with a linear gradient of imidazole mixed by (20 mM Tris-HCl, 2 M NaCl, 2 mM β-mercaptoethanol) and (20 mM Tris-HCl, 2 M NaCl, 2 mM β-mercaptoethanol, 500 mM imidazole). Peak fractions were collected and concentrated, and then loaded onto Superdex 200 10/300 GL (Cytiva) in buffer E (20 mM Tris-HCl pH 7.5, 2 M NaCl, 1 mM EDTA, 1 mM DTT). Peak fractions were analyzed by SDS-PAGE and fractions containing only histone octamer were pooled and concentrated and stored at −80 °C in small aliquots.

#### Nucleosome assembly

Nucleosome assembly was performed as previously described(*42, 58*) with some modifications. Briefly, the caps of 1.5-ml Eppendorf tube were used to make dialysis button. Histone octamer and deoxyinosine-containing Widom 601 DNA were mixed at a molar ratio of 1:1 in buffer E, and incubated for at least 0.5 hour on ice. The mixture was then transferred into dialysis buttons sealed with dialysis membrane. Dialysis buttons were transferred into dialysis bag filled with buffer E. The dialysis bag was dialyzed against buffer F (20 mM Tris-HCl pH 7.5, 50 mM NaCl, 1 mM EDTA, 1 mM DTT) for 17 hours at 4°C. To complete nucleosome core particles assembly, the dialysis buttons were dialyzed against fresh buffer F for additional 4 hours. Assembled nucleosome was verified by 6% TBE-PAGE and used for AAG-NCP complex assembly.

#### Preparation of AAG-NCP complex

All AAG-NCP complexes were prepared with the following method. Nucleosome was mixed with AAG protein at a molar ration of 1: 50 in buffer C. The complex was stabilized by GraFix(*59*) using TLS-55 rotor, the mixture was loaded onto a glycerol gradient of 20 mM HEPES pH 7.5, 100 mM NaCl, 0.1 mM EDTA, 1 mM DTT, 20% glycerol (v/v) and 20 mM HEPES pH 7.5, 100 mM NaCl, 0.1 mM EDTA, 1 mM DTT, 40% glycerol (v/v), 0.05% glutaraldehyde. The sample was centrifuged at 200,000g for 17 h at 4°C, and fractionated into 100 μl aliquots manually. 10 μl of 1 M Tris-HCl (pH 7.5) was added to each aliquot to quench crosslinking reaction. Each fraction was examined for AAG-NCP complex assembly by 6% TBE-PAGE. Fractions containing AAG-NCP complex were pooled and concentrated, and the glycerol removal and buffer exchange were achieved by ultrafiltration using 50k AmiconUltra-0.5 centrifugal filter unit. A final buffer of 20 mM Tris-HCl pH 7.5, 50 mM NaCl, 0.1 mM EDTA, 1 mM DTT was used for cryo-EM sample preparation.

### Cryo-EM sample preparation and data collection

AAG-NCP complexes concentrated to ∼1.0 mg/ml were used for cryo-grid preparation. Quantifoil holey carbon Au R1.2/1.3 grids were glow-discharged, and 4 μl of sample was applied to glow-discharged grids in Vitrobot Mark IV at 8°C and 100% humidity. After waiting for 10 s, the grids were plunged into the liquid ethane for vitrification. Grids were screened on a Talos Arctica (ThermoFisher) 200 kV TEM equipped with a K2 detector (Gatan). Data collection was performed with an FEI Titan Krios G2 TEM operated at 300 kV with a Gatan K2 (GIF) direct electron detector. Automated data collection was done using SerialEM(*60*). Data was collected with nominal magnification of ×130,000 (corresponding to a calibrated pixel size of 1.052 Å), with defocus range of −1.2 to −1.7 μm. For each movie stack, a total of 32 frames was collected at a dose rate of 8 e^-^ Å^-2^ s^-1^ for 8 s.

### Image processing

For each dataset, the movie stacks were first subjected to beam-induced motion correction, electron-dose weighting and two-fold binning using MotionCor2(*61*). Contrast transfer function (CTF) parameters of dose-weighted motion-corrected micrographs were estimated by program Gctf(*62*). Around 400 particles were manually picked to generate an initial set of 2D template for automatic particle picking. Two rounds of reference-free 2D classification (25 iterations) were applied after particle auto-picking and bad 2D classes were discarded. The initial 3D reference was produced using RELION3.1(*63*). All subsequent processing steps were performed using RELION 3.1 unless otherwise stated. The image processing workflow was summarized in Figures S2-S5.

For the AAG-NCP^-30^ dataset, a total of 3,814 raw micrographs were acquired and 1,421,601 particles were auto-picked. After 2D classification, 1,249,347 particles were kept and subjected to two rounds of 3D classification with no symmetry imposed, which generated two major states. One is a free NCP (189,350 particles), called Apo state. The other is the AAG bounded state (126,881 particles), called post-catalytic state (confirmed in the model building). The particles from these two states were re-extracted and re-centered using a box size of 200 pixels for 3D refinement. Two individual soft-edged masks were used for high-resolution refinement, leading to two density maps at resolutions of 3.0 Å (Apo state) and 3.1 Å (Post-catalytic state) (gold-standard Fourier shell correlation (FSC) 0.143 criteria). Application of CTF refinement and Bayesian polishing boosted the resolutions to 2.8 Å and 2.9 Å (Figure S2), respectively. To further improve the local density of the AAG region in the AAG bounded state, another round of mask-based 3D refinement was performed. Final density maps were corrected for the modulation transfer function (MTF) of the K2 Summit detector. The map sharpening was carried out by both RELION and DeepEMhancer(*64*). The B factors used in RELION were automatically estimated through the post-processing procedure and the local resolution maps were generated using ResMap(*65*) in RELION.

For the AAG-NCP^-50^ dataset, 1,583,682 particles were selected from 3,918 raw images. After cleaning up the dataset by two rounds of 2D classification and one round of 3D classification, 460,662 particles were left for further processing. For the Apo state, another round of 3D classification was applied and one class showing clear features was chosen and subjected to 3D refinement. Finally, the resolution of the Apo state was pushed to 2.9 Å. For the post-catalytic state, all particles from good classes of the first round 3D classification were merged and a small soft spherical mask centered at the extra density near SHL_-5_ of the nucleosome was generated to perform a focused 3D classification (skip alignment). A class (73,954 particles) exhibiting strong additional density was selected and further refined. The CTF refinement and Bayesian polishing procedures were subsequently applied to improve the resolution of the final map to 3.0 Å (Figure S3).

For the AAG-NCP^-53^ dataset, 4,374 raw images were obtained and 1,711,509 particles were auto-picked. The data processing steps of AAG-NCP^-53AP^ were similar to that of the AAG-NCP^-50^ dataset. A set of 129,507 and 98,691 particles were used to generate the final density map for the Apo state and AAG bound state at 2.8 Å and 3.1 Å, respectively (Figure S4).

For the AAG-NCP^-55^ dataset, 1,706,627 particles were auto-picked from 4,363 raw micrographs. Similarly, the final map of the Apo state was determined at 2.8-Å resolution. And the final map of the post-catalytic state was determined at 2.9-Å resolution (Figure S5).

### Model building and refinement

Cryo-EM structure of a canonical nucleosome (PDB code 7OHC)(*46*) was used as the initial template for modelling. For the NCP^-30I^, the crystal structure was first docked into the density map using UCSF ChimeraX(*66, 67*), and structures of histones and DNA were adjusted manually using Coot(*68*). The −30A was replaced with DI. For the AAG-NCP^-30AP^, the structures of the nucleosome (PDB code 7OHC) and AAG (PDB code 1BNK)(*36*) were similarly modelled, and the −30A was replaced with AP site based on local density. These atomic models were refined in real space using Phenix(*69*) with the geometry and secondary structure restraints applied. The refined structures were re-checked in Coot to adjust the side chain to proper locations. final atomic models were evaluated using Molprobity(*70*). Same modelling procedures were performed for other maps. The statistics of data collection and model validation were summarized in Table. 3.

### Structural analysis

The published NCP structure (EMDB-12900, PDB: 7OHC)(*46*) was used as a reference. Initial structural analysis indicated a scaling factor between the published map and our maps. Therefore, we further calibrated the voxel sizes of the published map and our maps using the crystal structure of a canonical NCP (PDB code 1KX5)(*71*) as a standard. Based on the calibration, the voxel size of the reference density map (EMDB-12900) should be 1.06 Å. Subsequently, the reference model (7OHC) was adjusted by real-space refinement against the calibrated reference map using Phenix. The resulting refined model was used as the atomic model of the canonical NCP for structural analysis. Structural comparisons were performed using UCSF ChimeraX, and atom models were aligned by histone H3 using the matchmaker function. RMSD of nucleosomal DNA was calculated based on atoms from DNA backbone (C1’, C2’, C3’, C4’, C5’, O3’, O4’, O5’, P, OP1, OP2). Buried surface area between nucleosomal DNA and octameric histone core was determined using buried area function in UCSF ChimeraX.

### Data availability

Atomic coordinates of the associated structures have been deposited to the Protein Data (PDB) with the following accession codes 7XFC (NCP^-30I^), 7XFH (AAG-NCP^-30AP^), 7XFI (NCP^-50I^), 7XFJ (AAG-NCP^-50AP^), 7XFL (NCP^-53I^), 7XFM (AAG-NCP^-53AP^), 7XFN (NCP^-55I^) and 7XNP (AAG-NCP^-55AP^), respectively. The corresponding maps have been deposited to the Electron Microscopy Data Bank (EMDB) under following accession numbers EMDB-33171 (NCP^-30I^), EMDB-33172 (AAG-NCP^-30AP^), EMDB-33173 (NCP^-50I^), EMDB-33174 (AAG-NCP^-50AP^), EMDB-33175 (NCP^-53I^), EMDB-33176 (AAG-NCP^-53AP^), EMDB-33177 (NCP^-55I^) and EMDB-33322 (AAG-NCP^-55AP^), respectively. All data is available from the corresponding author.

## Acknowledgement

We thank Prof. Li Qing (Peking University) for sharing plasmids of *Xenopus laevis* histones. We thank the Core Facilities of Peking University School of Life Sciences for assistance with negative-staining electron microscopy; the Cryo-EM Platform and the Electron Microscopy Laboratory of Peking University for cryo-EM data collection; the High-performance Computing Platform of Peking University for help with computation. This work was supported by the National Science Foundation of China (31725007 to N.G.), the Ministry of Science and Technology of China (2019YFA0508904 to N.G.), and the Qidong-SLS Innovation Fund to N.G.

## Author Contributions

N.G. supervised the project. B.T. conceived the project; B.T. and L.Z. designed experiments; L.Z. and B.T. performed all experiments; L.Z. and B.T. performed EM analysis; B.T., L.Z. and N.G. wrote the manuscript.

## Competing interests

The authors declare no competing interests.

---

**Figure S1.**
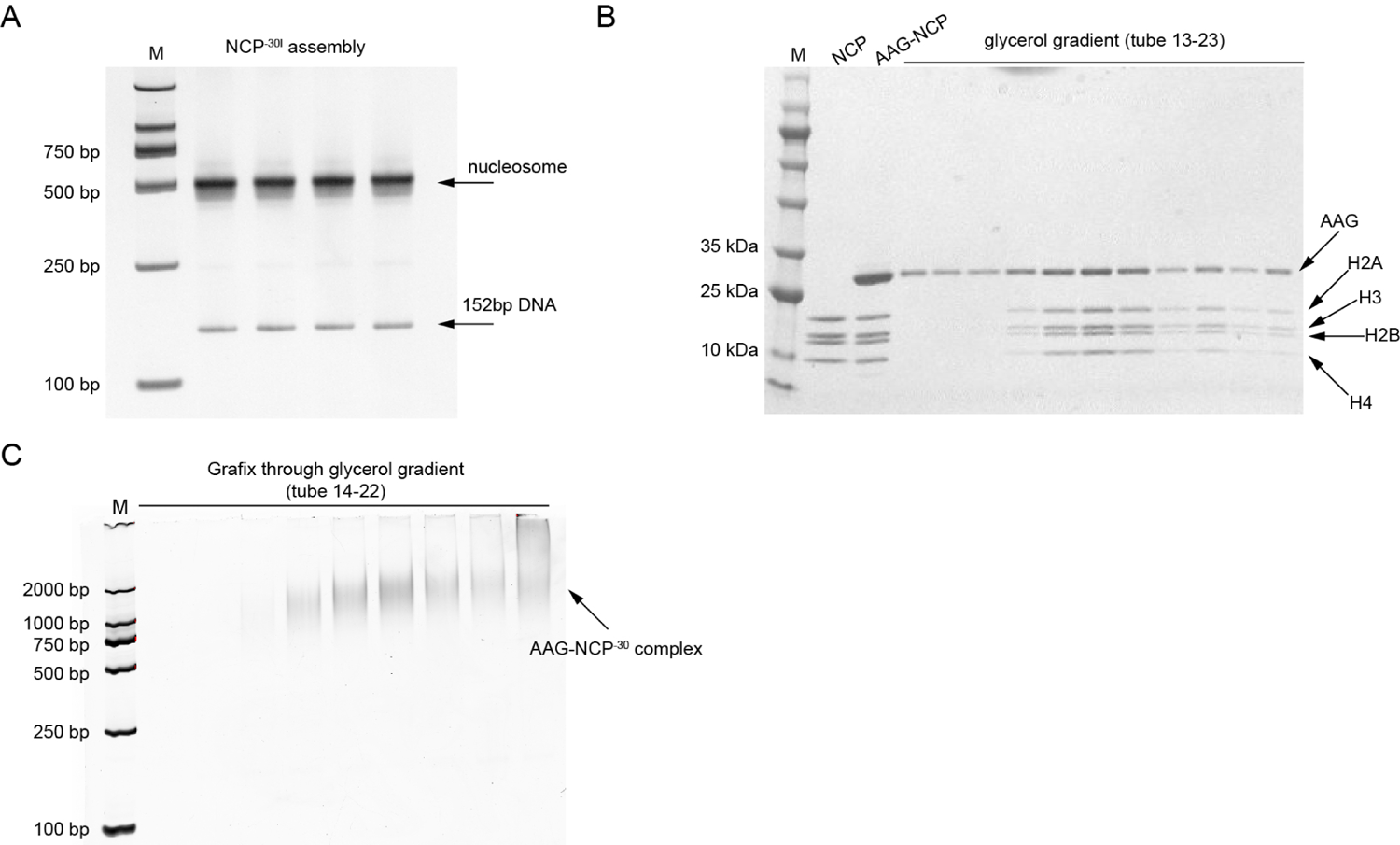
Assembly of the AAG-NCP^-30^ complex. **A,** TBE-PAGE (6%) analysis of the reconstituted nucleosomes bearing a deoxyinosine at −30. The four lanes represent four independent assembly reactions. **B,** SDS-PAGE analysis (4-20%) of the glycerol gradient fractions containing the AAG-NCP complexes. NCP, reconstituted NCP complexes; AAG-NCP, the reaction mixture. **C,** TBE-PAGE analysis (6%) of the GraFix fractions containing the AAG-NCP^-30^ complexes.

**Figure S2.**
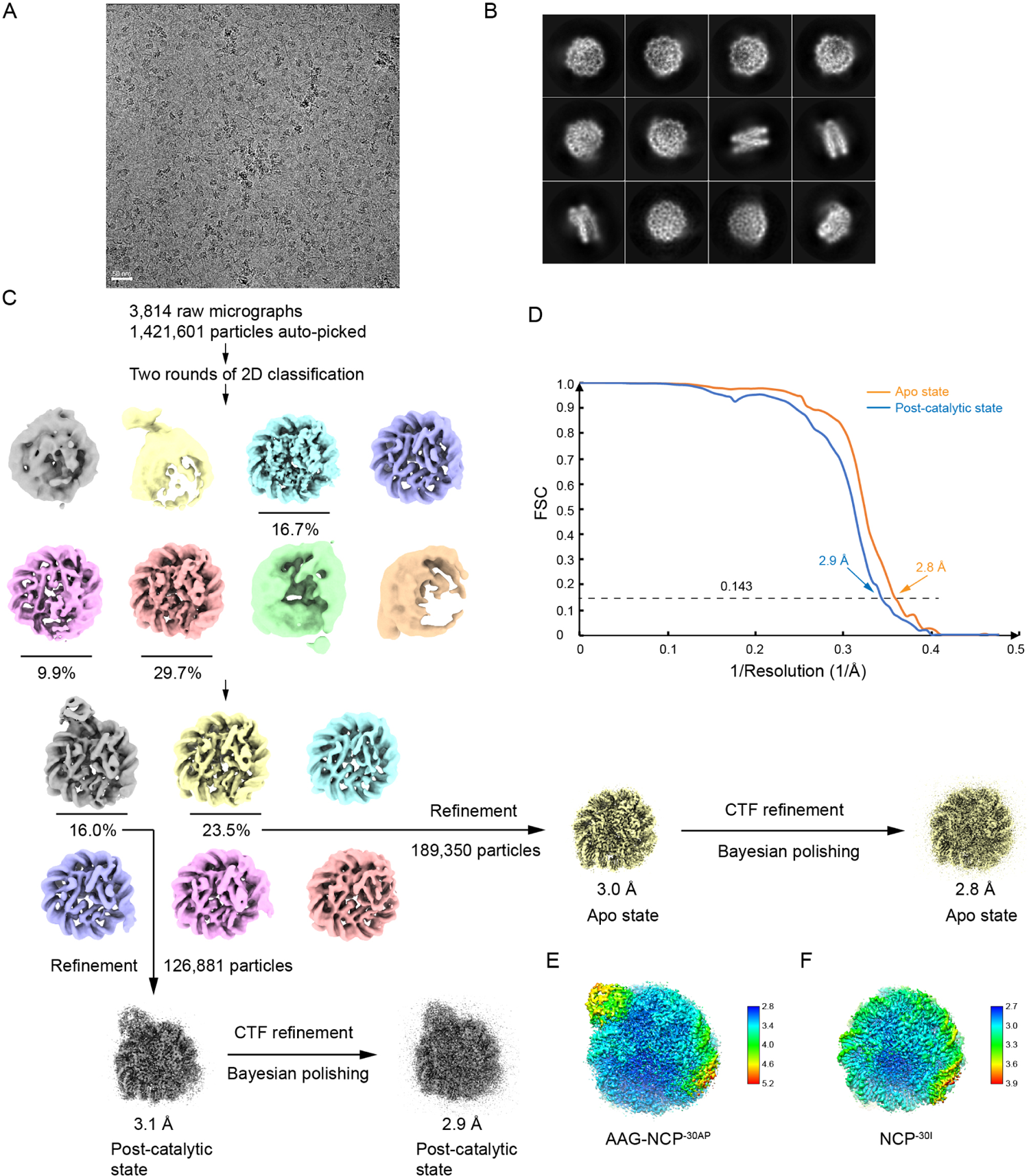
Data processing of the AAG-NCP^-30^ dataset. **A,** A representative raw cryo-EM image of the AAG-NCP^-30^ sample. **B,** Representative 2D classes of the AAG-NCP^-30^ particles. **C,** Image-processing workflow of the AAG-NCP^-30^ dataset. **D,** Gold-standard FSC curves of the final cryo-EM maps. **E,** Final local resolution estimation of the AAG-NCP^-30AP^ map. **F,** Final local resolution estimation of the NCP^-30I^ map.

**Figure S3.**
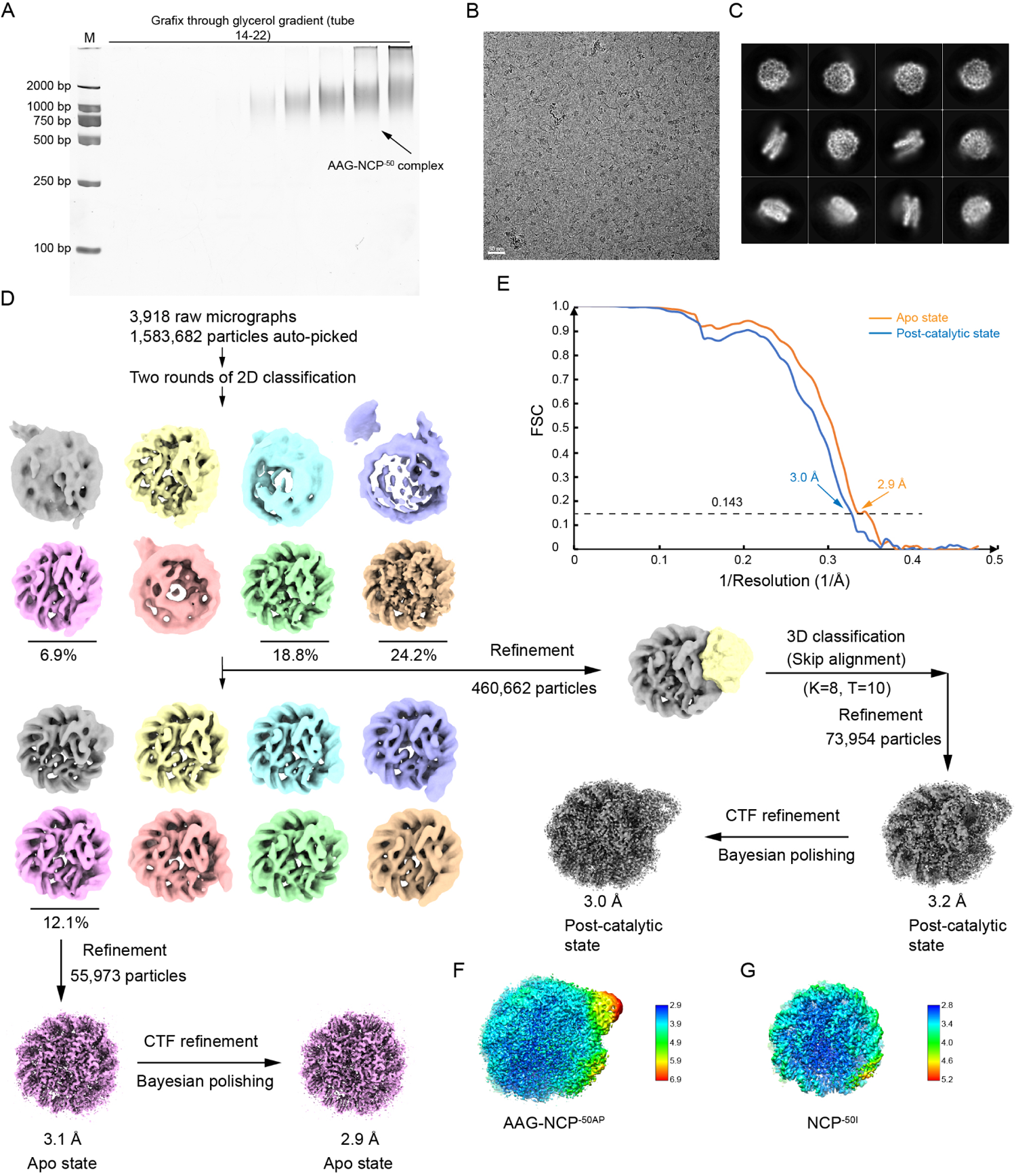
Data processing of AAG-NCP^-50^ dataset. **A,** TBE-PAGE analysis (6%) of the GraFix fractions containing the AAG-NCP^-50^ complexes. **B,** A representative raw cryo-EM image of the AAG-NCP^-50^ sample. **C,** Representative 2D classes of the AAG-NCP^-50^ particles. **D,** Image-processing workflow of the AAG-NCP^-50^ dataset. **E,** Gold-standard FSC curves of the final cryo-EM maps. **F,** Final local resolution estimation of the AAG-NCP^-50AP^ map. **G,** Final local resolution estimation of the NCP^-50I^ map.

**Figure S4.**
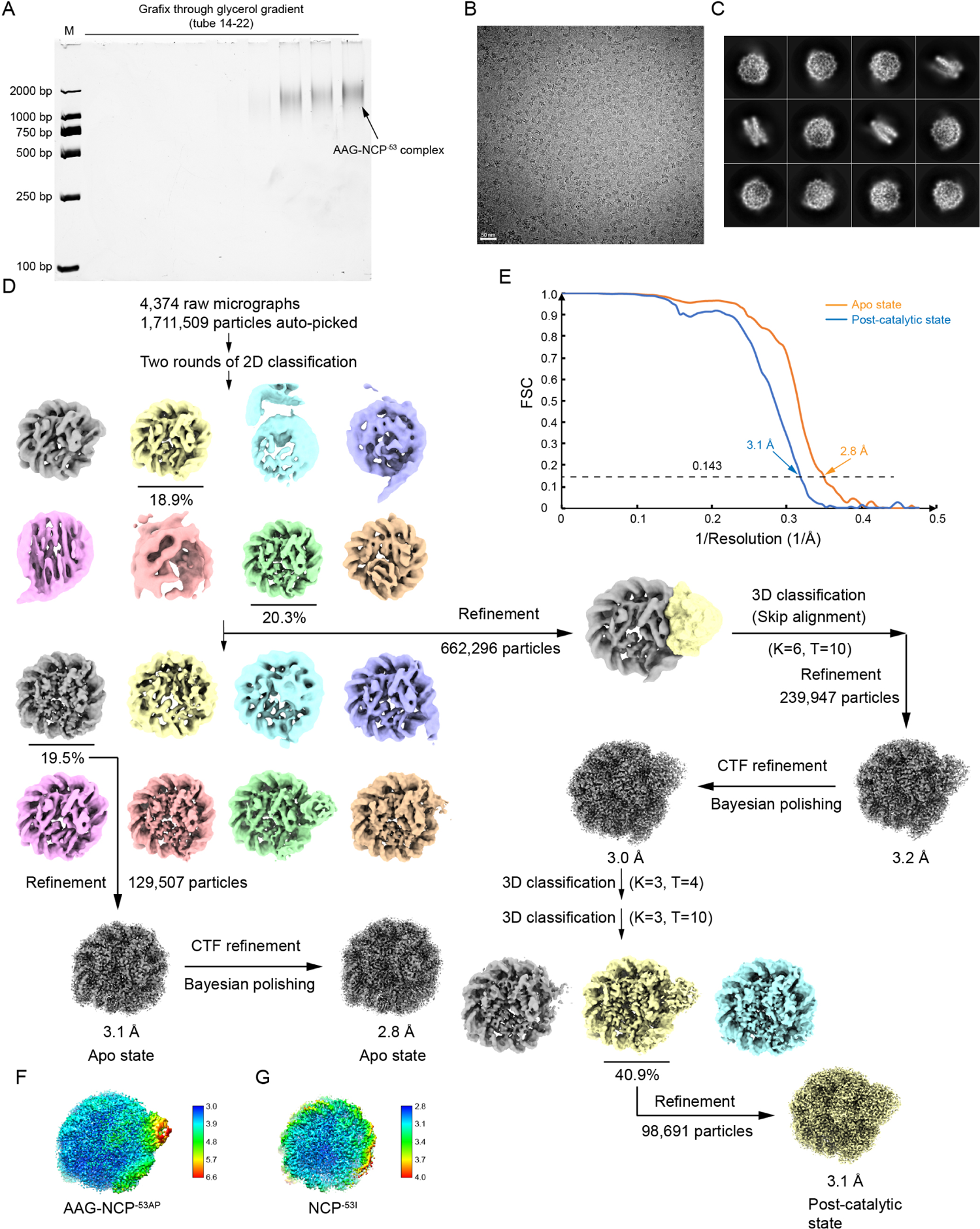
Data processing of AAG-NCP^-53^ dataset. **A,** TBE-PAGE analysis (6%) of the GraFix fractions containing the AAG-NCP^-53^ complexes. **B,** A representative raw cryo-EM image of the AAG-NCP^-53^ sample. **C,** Representative 2D classes of the AAG-NCP^-53^ particles. **D,** Image-processing workflow of the AAG-NCP^-53^ dataset. **E,** Gold-standard FSC curves of the final cryo-EM maps. **F,** Final local resolution estimation of the AAG-NCP^-53AP^ map. **G,** Final local resolution estimation of the NCP^-53I^ map.

**Figure S5.**
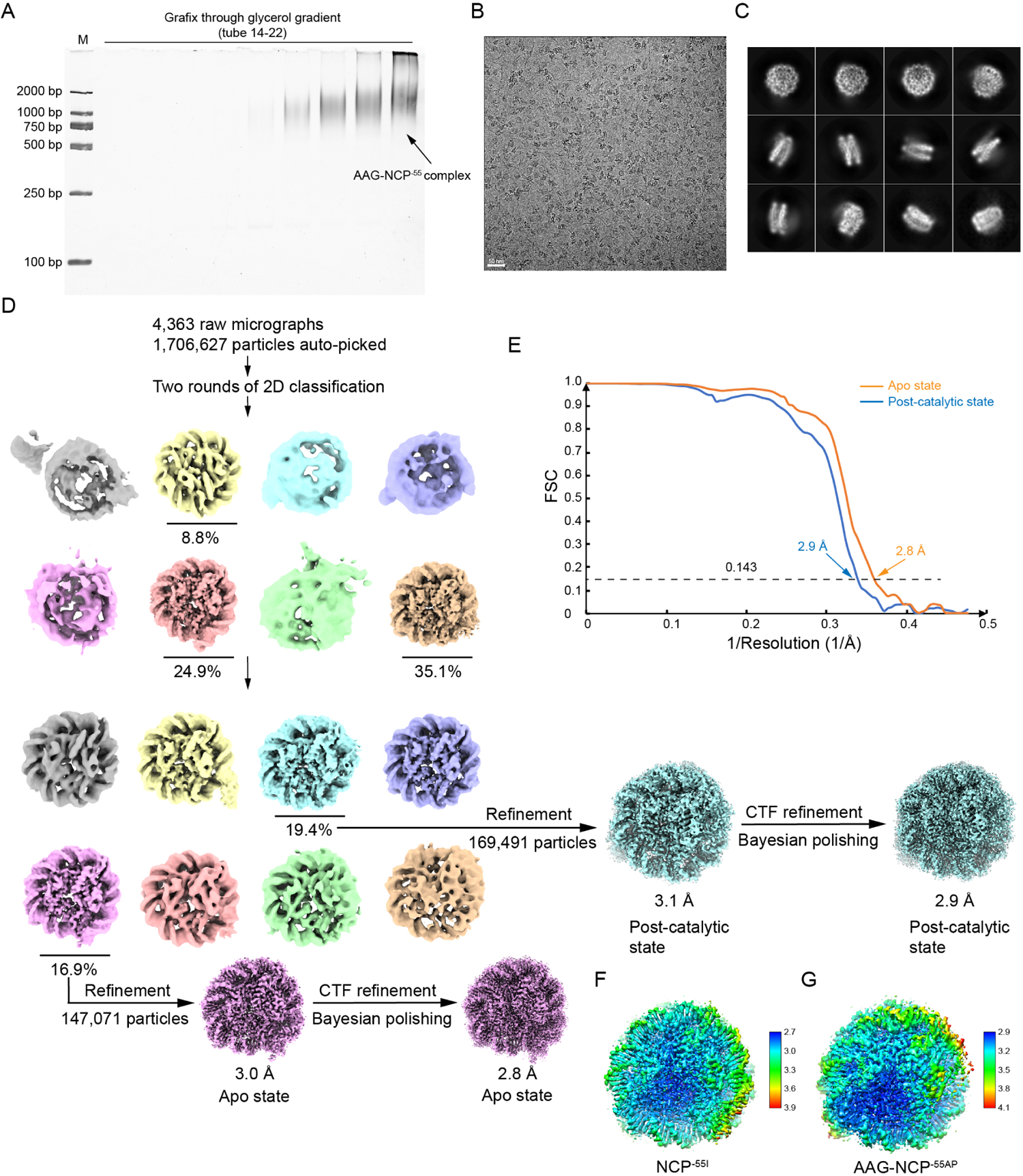
Data processing of AAG-NCP^-55^ dataset. **A,** TBE-PAGE analysis (6%) of the GraFix fractions containing the AAG-NCP^-55^ complexes. **B,** A representative raw cryo-EM image of the AAG-NCP^-55^ sample. **C,** Representative 2D classes of the AAG-NCP^-55^ particles. **D,** Image-processing workflow of the AAG-NCP^-55^ dataset. **E,** Gold-standard FSC curve of the final cryo-EM maps. **F,** Final local resolution estimation of the NCP^-55I^ map. **G,** Final local resolution estimation of the AAG-NCP^-55AP^ map.

**Figure S6.**
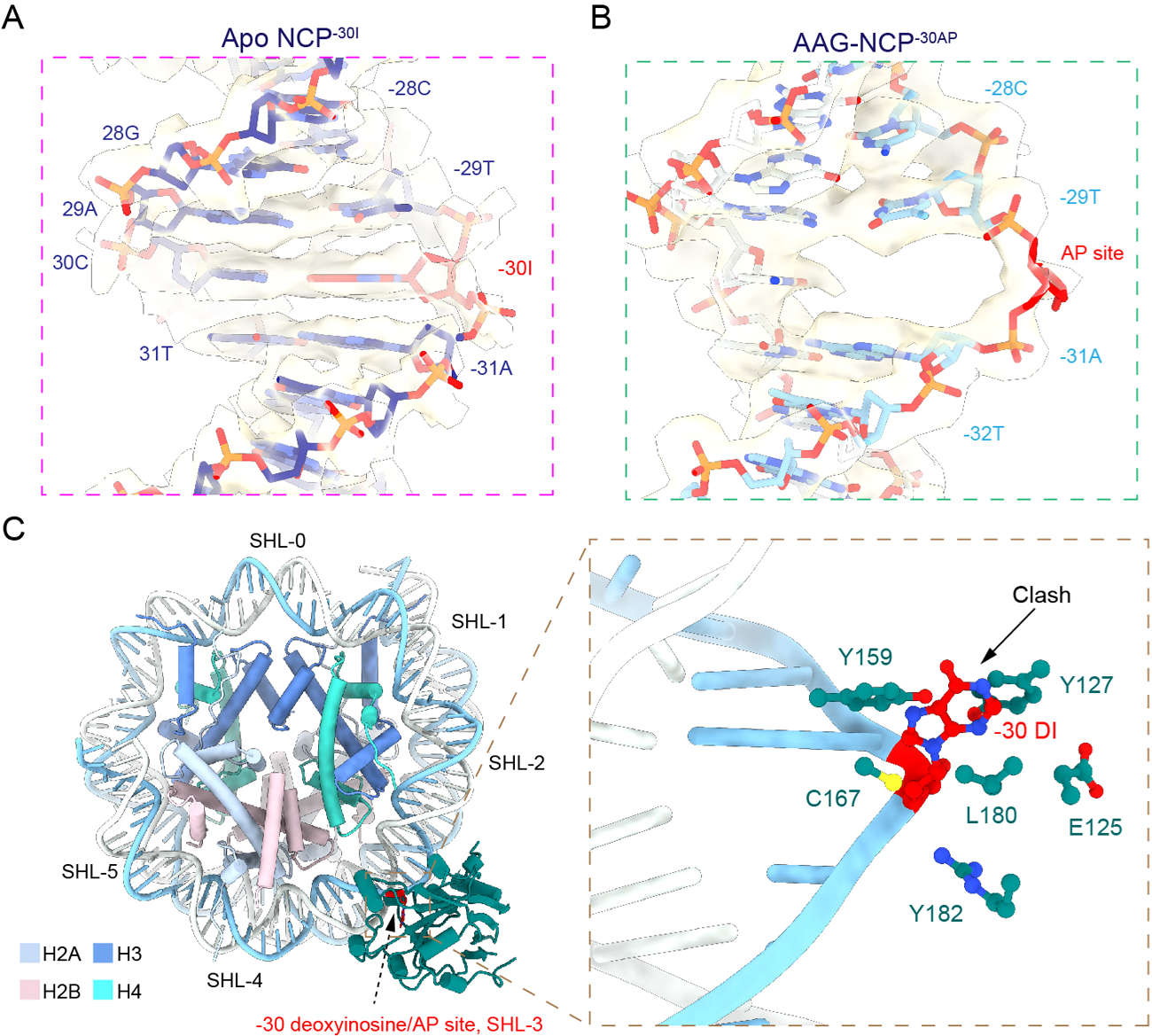
Local density of the NCP^-30I^ and AAG-NCP^-30AP^ maps around the damaged base. **A,** Local cryo-EM density map of the nucleosomal DNA around deoxyinosine at −30 in the NCP^-30I^. **B,** Local cryo-EM density map of the nucleosomal DNA around the AP-site at −30 in the AAG-NCP^-30AP^. **C,** Superimposition of a deoxyinosine onto the model of the AAG-NCP^-30AP^, highlighting the steric clash between the DI and AAG residues.

**Figure S7.**
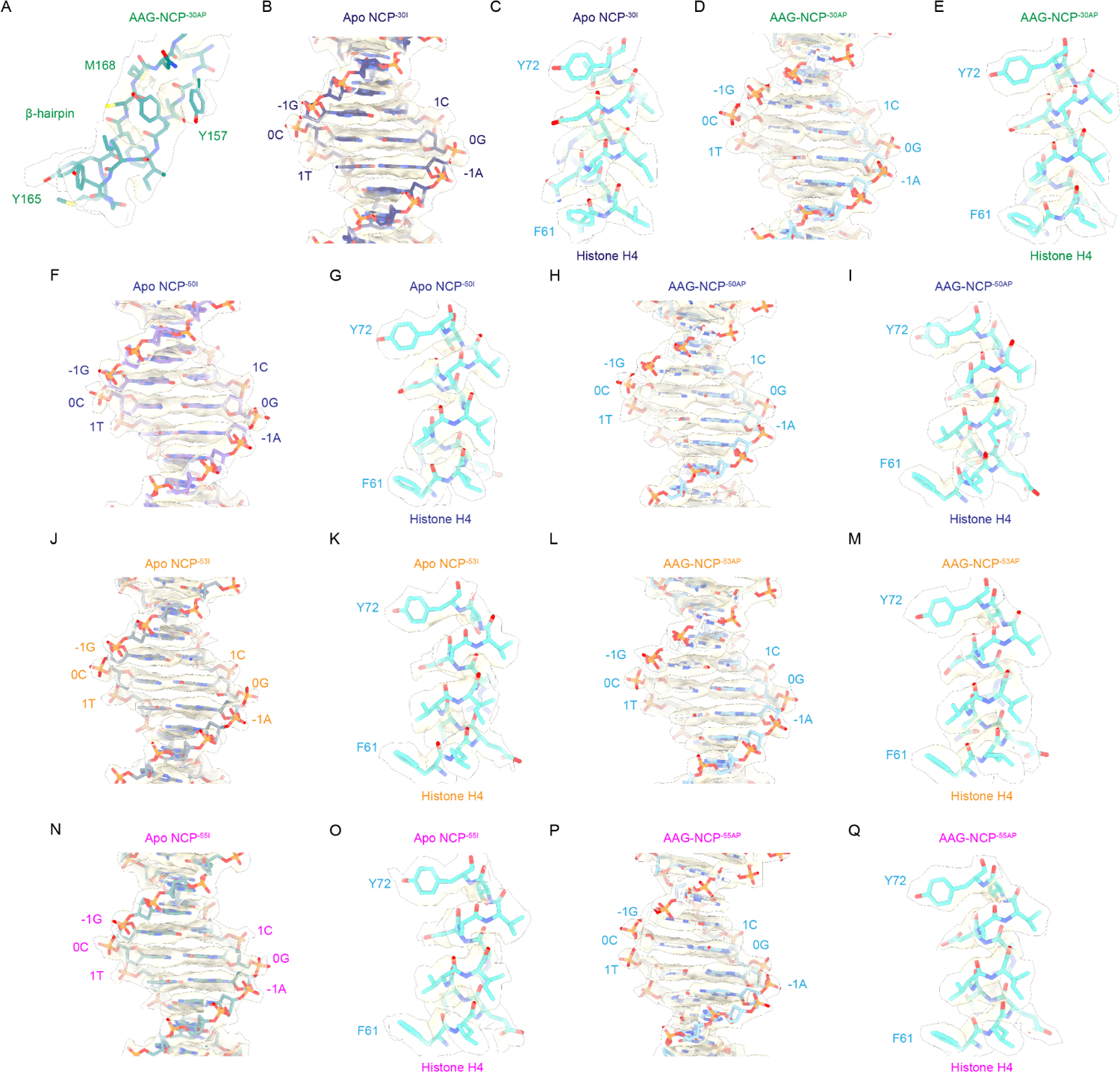
Evaluation of cryo-EM density maps. **A-Q**. Representative local cryo-EM density maps of AAG, nucleosomal DNA and Histone H4 for each cryo-EM map.

**Figure S8.**
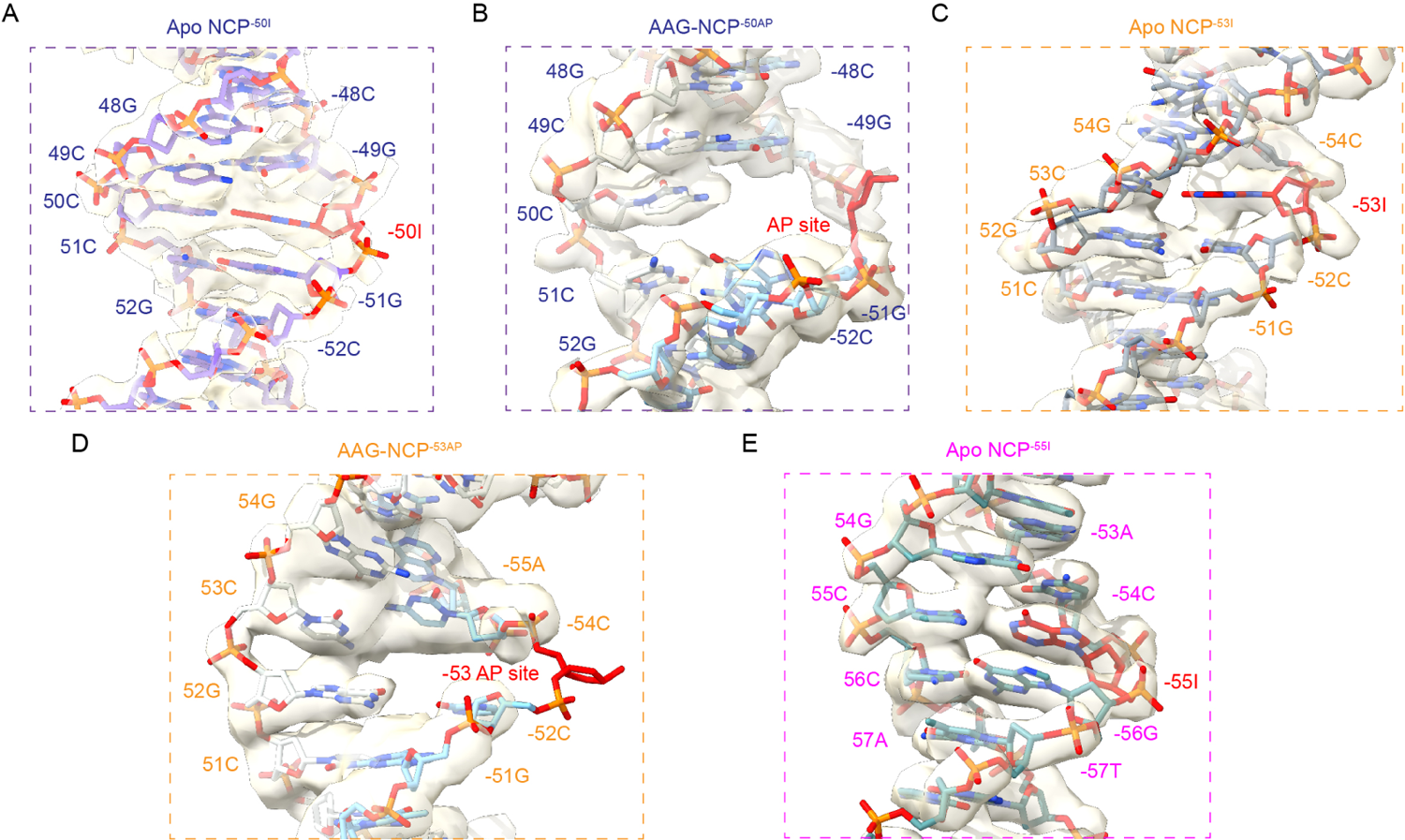
Local density of the NCP^-50I^, NCP^-53I^, NCP^-55I^, AAG-NCP^-50AP^, and AAG-NCP^-53AP^ maps in the region of the damaged base. **A-B,** Local cryo-EM density map of the nucleosomal DNA around −50 in the NCP^-50I^ and AAG-NCP^-50AP^. **C-D**, Local cryo-EM density map of the nucleosomal DNA around −53 in the NCP^-53I^ and AAG-NCP^-53AP^. **E**, Local cryo-EM density map of the nucleosomal DNA around −55 in the NCP^-55I^.

**Figure S9.**
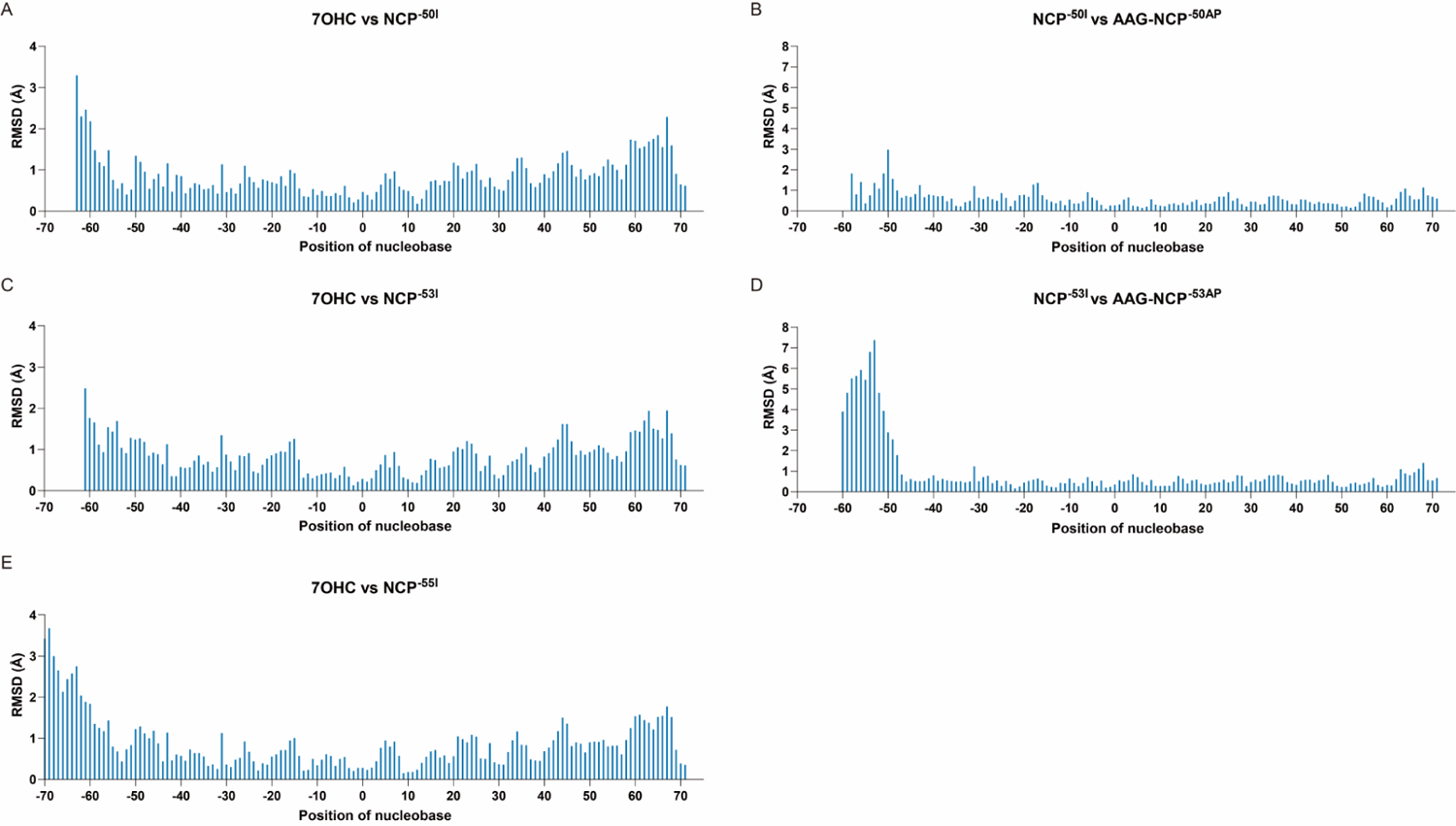
RMSD plots for pairwise comparisons of the nucleosomal DNA distortion. **A-B,** RMSD of the DNA backbone between a canonical NCP (PDB: 7OHC) and the NCP^-50I^, and between the NCP^-50I^ and the AAG-NCP^-50AP^. **C-D,** RMSD of the DNA backbone between a canonical NCP (PDB: 7OHC) and the NCP^-53I^, and between the NCP^-53I^ and the AAG-NCP^-53AP^. **E,** RMSD of the DNA backbone between a canonical NCP (PDB: 7OHC) and the NCP^-55I^. The RMSD of each residue is plotted against residue number. Note that the terminal DNA (from ∼60 to exit) of the NCP^-50I^, AAG-NCP^-50AP^, NCP^-53I^ and AAG-NCP^-53AP^ is relatively flexible in the maps and not modelled, and therefore not included for RMSD calculation.

**Figure S10.**
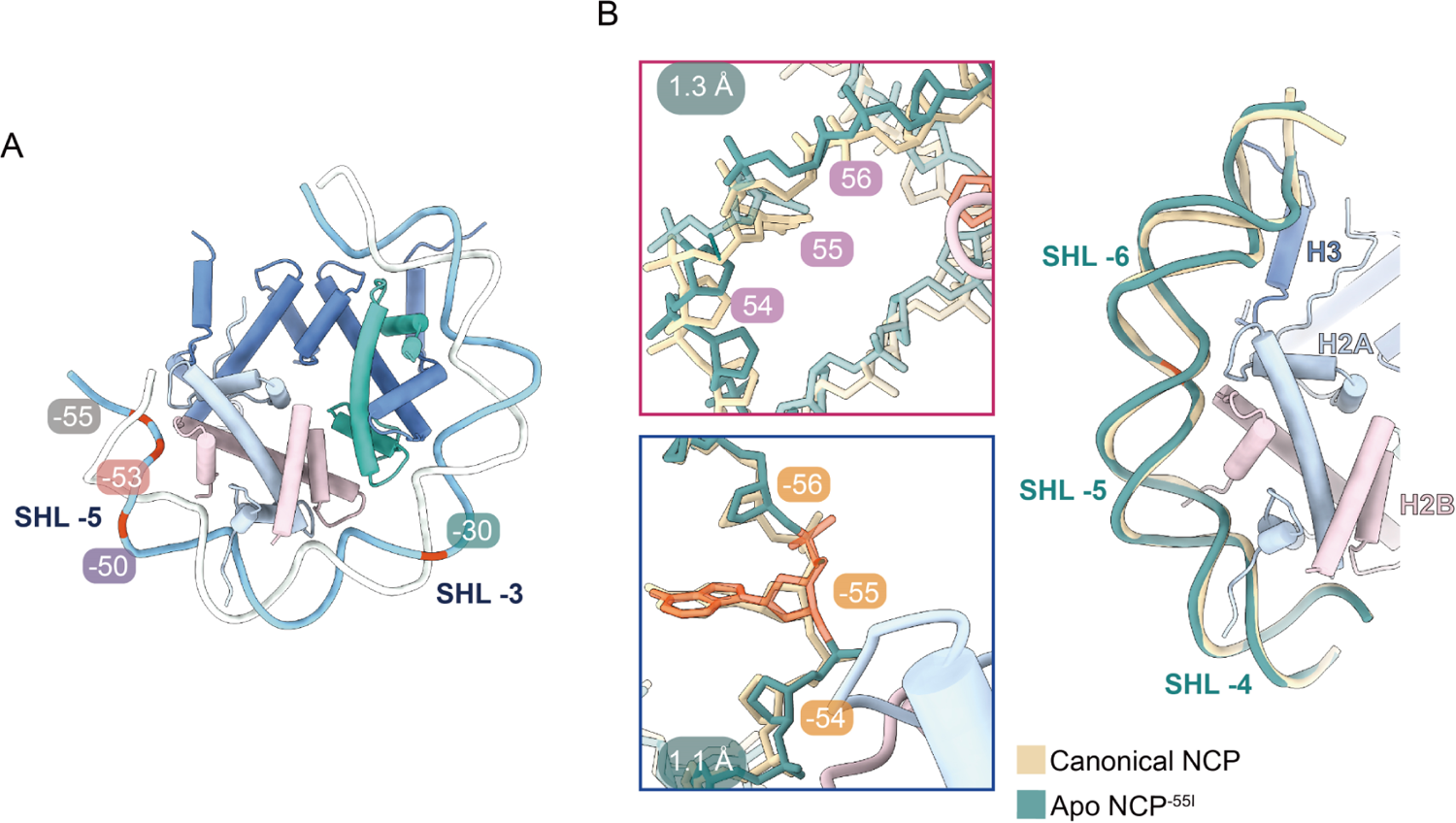
Nucleosomal DNA deformation in the NCP^-55I^. **A,** The position of −55 in the nucleosome. **B,** The nucleosomal DNA perturbation of the NCP^-55I^ in comparison with a canonical NCP (PDB: 7OHC). RMSD of the DNA backbone is 1.3 Å for the damaged strand from 50 to 60, and 1.1 Å for the undamaged strand from −50 to −60.

**Figure S11.**
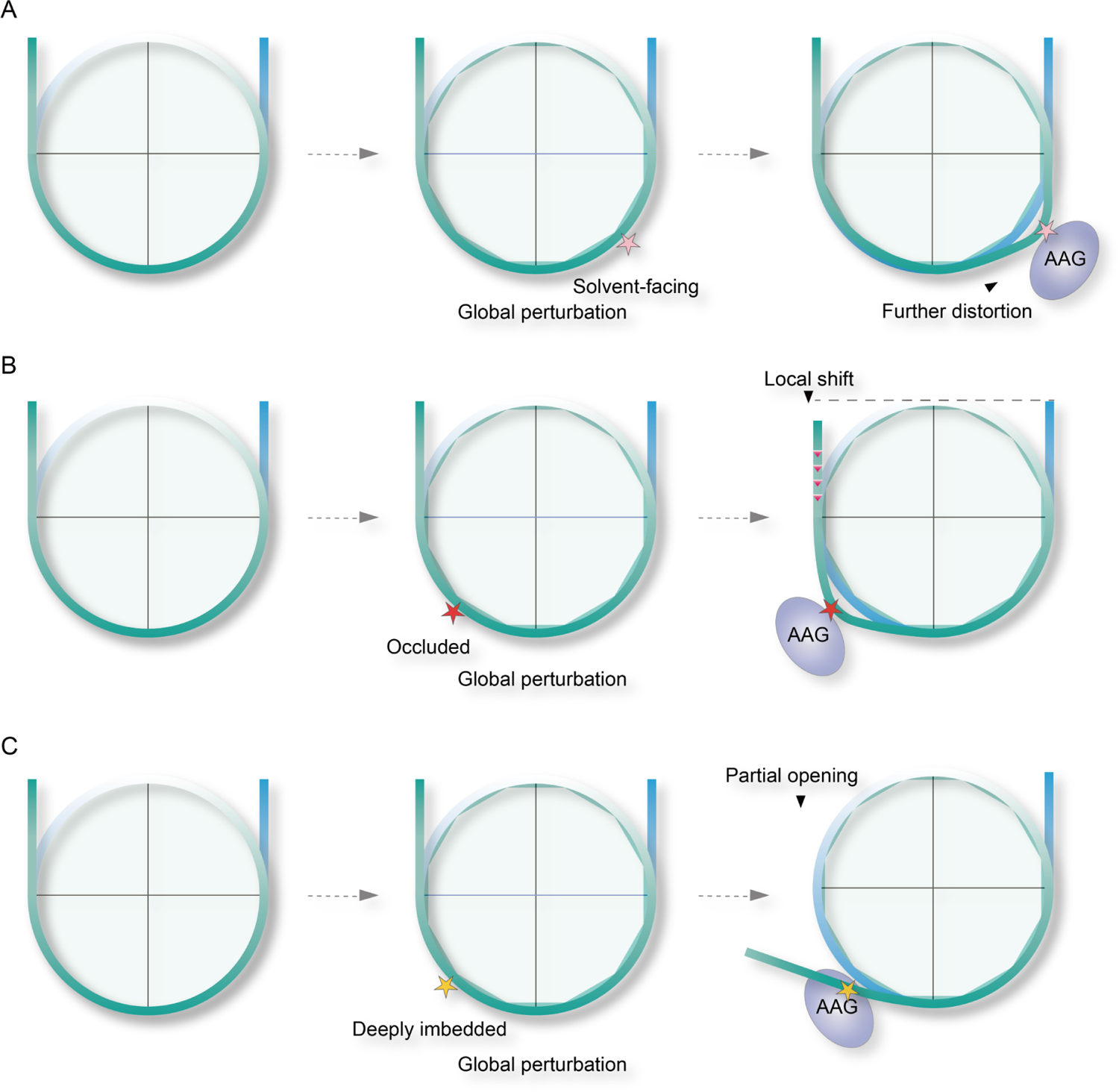
Models of AAG-mediated base excision in nucleosome. **A,** For solvent-facing position with high solution accessibility, by forming a stable AAG-NCP complex, AAG augments local DNA distortion to fulfill its function. **B,** For occluded position with medium solution accessibility, the binding of AAG imposes drastic local DNA distortion and causes additional local register shift of DNA to avoid clash between flipped base and histones. **C,** For deeply embedded position with low solution accessibility, local DNA distortion and translocation are insufficient to expose the damaged site, the global perturbation caused by deoxyinosine increases the possibility of spontaneous DNA unwrapping. AAG captures the damaged base in peeled-off DNA, and forms a stable AAG-NCP complex.

**Movie. 1.** Local DNA distortion caused by AAG engagement in AAG-NCP^-50AP^. Comparison between NCP^-50I^ and AAG-NCP^-50AP^.

**Movie. 2.** Close-up view of local DNA distortion caused by AAG engagement in AAG-NCP^-50AP^. Close-up view of DNA distortion around −50 when comparing NCP^-50I^ to AAG-NCP^-50AP^.

**Movie. 3.** Local DNA distortion caused by AAG engagement in AAG-NCP^-53AP^. Comparison between NCP^-53I^ and AAG-NCP^-53AP^.

**Movie. 4.** Close-up view of local DNA distortion caused by AAG engagement in AAG-NCP^-53AP^. Close-up view of DNA distortion around −53 when comparing NCP^-53I^ to AAG-NCP^-53AP^.

## Notes

### Competing Interest Statement

The authors have declared no competing interest.

